# Variation in Base Composition Underlies Functional and Evolutionary Divergence in Non-LTR Retrotransposons

**DOI:** 10.1101/788562

**Authors:** Robert P. Ruggiero, Stéphane Boissinot

## Abstract

**Background:** Non-LTR retrotransposons often exhibit base composition that is markedly different from the nucleotide content of their host’s gene. For instance, the mammalian L1 element is AT-rich with a strong A bias on the positive strand, which results in a reduced transcription. It is plausible that the A-richness of mammalian L1 is a self-regulatory mechanism reflecting a trade-off between transposition efficiency and the deleterious effect of L1 on its host. We examined if the A-richness of L1 is a general feature of non-LTR retrotransposons or if different clades of elements have evolved different nucleotide content. We also investigated if elements belonging to the same clade evolved towards different base composition in different genomes or if elements from the same clades evolved towards similar base composition in the same genome.

**Results:** We found that non-LTR retrotransposons differ in base composition among clades within the same host but also that elements belonging to the same clade differ in base composition among hosts. We showed that nucleotide content remains constant within the same host over extended period of evolutionary time, despite mutational patterns that should drive nucleotide content away from the observed base composition.

**Conclusions:** Our results suggest that base composition is evolving under selection and may be reflective of the long-term co-evolution between non-LTR retrotransposons and their host. Finally, the coexistence of elements with drastically different base composition suggests that these elements may be using different strategies to persist and multiply in the genome of their host.

## BACKGROUND

Non-LTR retrotransposons (nLTR-RTs) are ubiquitous in vertebrate genomes and have profoundly affected the size, structure and function of these genomes [1–4]. nLTR-RTs constitute a diverse and ancient group of transposable elements whose origin predates the diversification of the main eukaryotic lineages [5]. They can be classified into 25 clades that differ in the number of open-reading frames (ORFs - one or two) and the presence of functional motifs [6]. The mode of mobilization of nLTR-RTs has not been elucidated for all clades but it is likely that, considering their structural similarities, all these elements transpose via a target-primed reverse transcription reaction, as experimentally demonstrated for the R2Bm and L1 elements [7, 8].

Since nLTR-RTs are rarely transmitted horizontally in vertebrates (with the exception of elements of the RTE clade [9–11]), the interaction between nLTR-RTs and the genome of their host is among the most intimate and long-lasting co-evolutionary processes found in nature. nLTR-RTs have been a source of evolutionary novelties [3], yet they can also be a source of deleterious alleles [12–14]. Thus, vertebrate hosts have evolved a number of repression mechanisms that have in turn affected the evolution of the nLTR-RT [15]. Vertebrate genomes and nLTR-RTs have shaped each other’s evolutionary fate and the signature of these interactions can be seen in the sequence of retrotransposons. For instance, the ORF1 of mammalian L1 carry the signature of adaptive evolution [16–18], whereas the recurrent replacement of the promoter region during L1 evolution is indicative of an arms race between L1 and host-encoded repressors of transcription [16, 17, 19, 20].

The base composition of nLTR-RTs may also reflect the nature of the interactions between elements and their hosts. The L1 sequence in mammals is AT-rich with an A bias on the positive strand [16]. From the perspective of L1 transposition, this base composition is not optimal since A-rich sequences are poorly transcribed, due, in part, to the presence of premature poly-adenylation signals causing early termination of transcription [21, 22]. Indeed, a synthetic codon-optimized L1 element replicates much more effectively in a retrotransposition assay [23]. In addition, the A-richness of L1 makes the codon usage of its ORFs poorly adapted for efficient translation [24]. This raises the possibility that the unusual base composition of L1 is a mechanism of self-regulation [25] and may reflect a trade-off between transposition efficiency and the deleterious effect of L1 on its host [21]. For instance, an L1 element with an unbiased base composition will replicate more effectively but this could result in a rate of transposition that is so deleterious for the host that such an element would not persist in the population.

The base composition is one of the most fundamental properties of a DNA sequence because it profoundly affects a number of important functions such as the efficacy of transcription [22, 26], the secondary structure of DNA and RNA molecules, the codon usage [27, 28] as well as the amino acid composition of encoded proteins [29]. All these aspects can potentially affect the reaction of retrotransposition and the overall replicative success of the element [22]. In addition, the base composition of an element can affect the host upon insertion by decreasing the efficacy of transcription of host genes [30], which explains the rarity of AT-rich L1 elements in introns [31], or by modifying the epigenetic status of the region where it inserts [32], for instance by providing novel CpG sites in the case of a GC-rich element. However, since the pioneering work of Lerat et al. [24, 33], the evolution of base composition in transposable elements has not been analyzed in detail, although the number of available genome sequences has drastically increased since these early studies. Here we performed an analysis of the base composition of nLTR-RT in vertebrates. Our goal was to determine if the A-richness of mammalian L1 is a general feature of nLTR-RT or if different clades of nLTR-RT have evolved different nucleotide composition. We also tested if there is a host effect, so that elements belonging to the same clade evolved towards different base composition in different genomes, or if elements from different clades tend to evolve toward similar base composition when in the same genome.

## RESULTS

We examined the evolution of base composition in the major clades of nLTR-RT represented in vertebrates (L1, Tx1, RTE, I, Rex1, CR1, L2, R4 and Penelope – the entire dataset is available as supplementary material 1). Our dataset consists of 193 consensus sequences which correspond to the most recently active families (<5% divergence among sequences within family) in 14 species of mammals (cow, pig, horse, rabbit, human, lemur, armadillo, dog, panda, hyrax, elephant, rat, mouse and opossum), a reptile (the green anole *Anolis carolinensis*), an amphibian (the African clawed frog *Xenopus tropicalis*) and five teleost fish (the zebrafish *Danio rerio*, the three-spined stickleback *Gasterosteus aculeatus*, the medaka *Oryzias latipes*, the fugu *Takifugu rubripes* and the pufferfish *Tetraodon nigroviridis*). The number of sequences, and the diversity of clades differed considerably among organisms, from one consensus sequence in each mammalian species, all belonging to the L1 clade, to 56 in the frog, where 6 clades are represented. The genomes analyzed here contain much larger numbers of nLTR-RT families, but we chose to limit our analysis to recently active families to avoid uncertainties when constructing consensus sequences. Figure 1 shows the phylogenetic relationships among the sequences analyzed here based on the reverse-transcriptase domain of ORF2. The major recognized clades of nLTR-RTs were recovered with good statistical support, although relationships among clades are not well supported. The main exception is the CR1 clade, which appears paraphyletic. However, further analyses based on the entire ORF2 support the reciprocal monophyly of the CR1 and L2 clades (see below).

**Figure 1.**
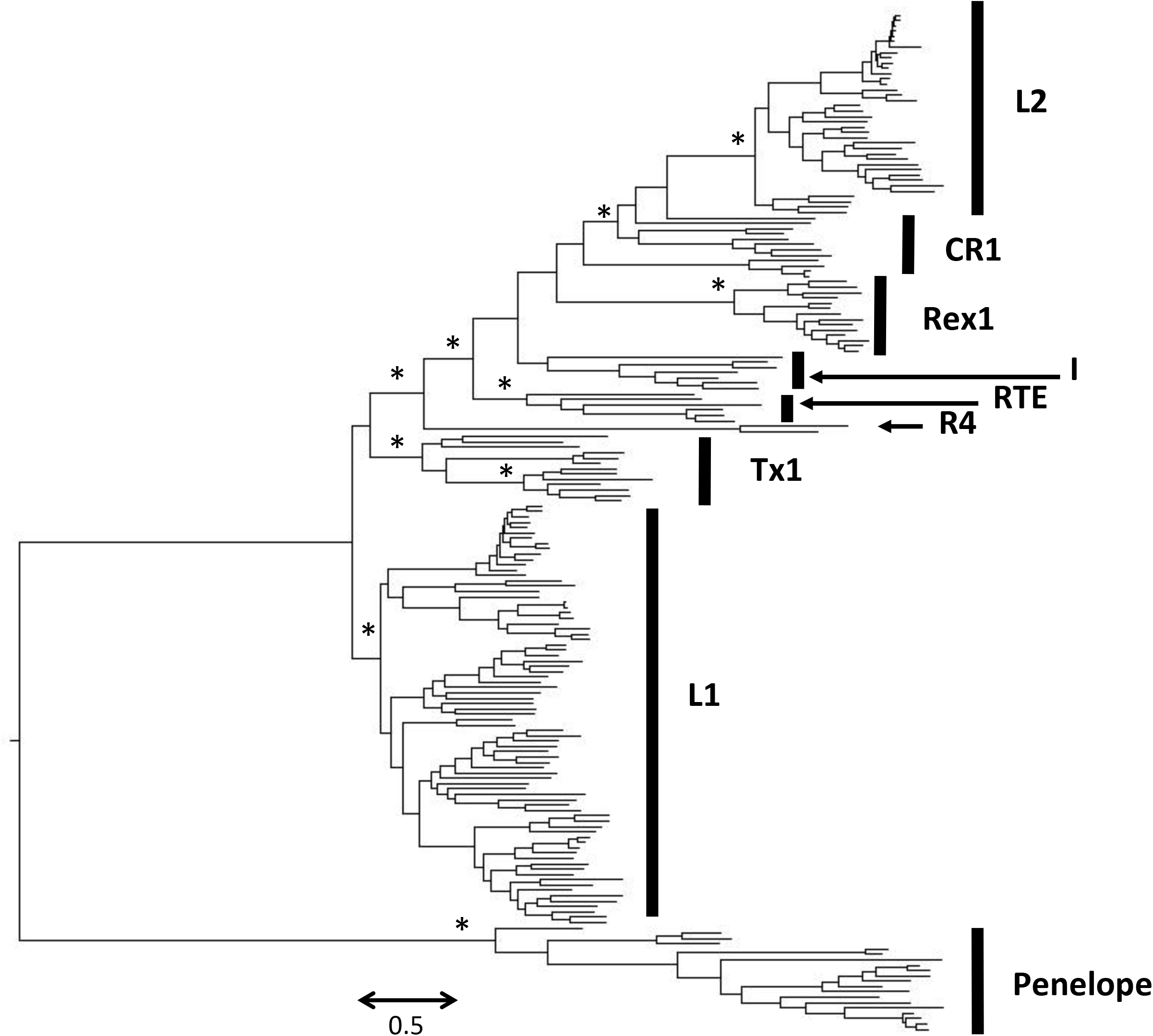
Phylogenetic relationships among nLTR-RT retrotransposons based on an amino-acid alignment of the reverse-transcriptase domain. The tree was built with the maximum likelihood method using the LG+G model of mutation and its robustness was assessed by 500 bootstrap replicates. Nodes that are supported by bootstrap values higher than 70% are indicated with an asterisk.

### Base composition varies among vertebrate lineages and among nLTR-RT clades

We first compared the base composition of the ORF that encodes the reverse transcriptase activity (homologous to the human L1 ORF2) since it is the only region that is shared among all clades of nLTR-RT and because additional ORFs, when present, are not homologous among clades. We first performed a comparison at the level of vertebrate class (mammals, reptiles, amphibians and teleost fish; figure 2 and table 1).

**Figure 2.**
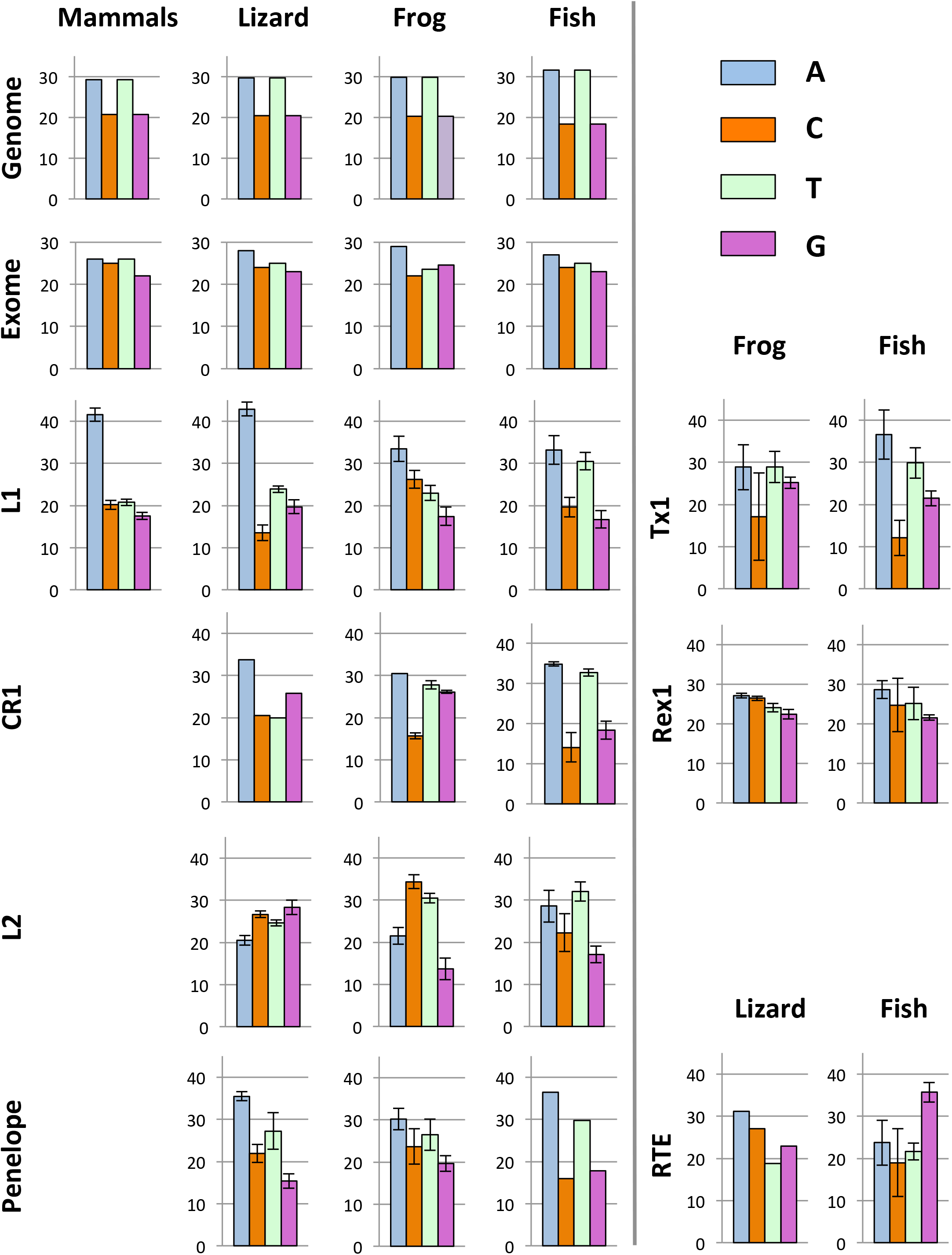
Base composition of ORF2 for the major clades of nLTR-RT in mammals, lizard, frog and teleostean fish. For comparison, the base composition of the host’s genomes and exomes are indicated.

**Table 1.** Statistics describing the base composition of ORF2 in nLTR-RT.

| Clades | Organisms | N | %AT | %GC | GC1 | GC2 | GC3 | PolyA | Nc | CAI | eCAI | RCDI | eRCDI |
| --- | --- | --- | --- | --- | --- | --- | --- | --- | --- | --- | --- | --- | --- |
| L1 | Mammals | 14 | 62.3 ± 1.3 | 37.7 ± 1.3 | 39.5 ± 1.2 | 32.9 ± 1.0 | 40.7 ± 2.2 | 13.5 ± 3.5 | 45.3 | 0.74 | 0.77 | 1.41 | 1.54 |
|  | Lizard | 12 | 66.8 ± 1.1 | 33.2 ± 1.1 | 37.2 ± 1.4 | 29.4 ± 0.8 | 33.2 ± 2.4 | 23.0 ± 6.5 | 44.8 | 0.72 | 0.75 | 1.65 | 1.71 |
|  | Frog | 35 | 56.4 ± 3.1 | 43.6 ± 3.1 | 47.4 ± 3.3 | 38.3 ± 2.2 | 42.8 ± 4.5 | 6.3 ± 3.5 | 50.9 | 0.79 | 0.84 | 1.25 | 1.306 |
|  | Fish | 18 | 63.7 ± 2.1 | 36.3 ± 2.1 | 39.6 ± 2.4 | 30.6 ± 2.2 | 32.1 ± 2.8 | 11.7 ± 7.4 | 43.1 | 0.72 | 0.75 | 1.71 | 1.92 |
| Tx1 | frog | 2 | 57.7 ± 9.0 | 42.3 ± 9.0 | 48.0 ± 9.2 | 35.7 ± 7.2 | 43.1 ± 10.6 | 6.0 ± 5.7 | 50.3 | 0.84 | 0.84 | 1.20 | 1.38 |
|  | fish | 11 | 61.4 ± 9.6 | 38.6 ± 9.6 | 44.6 ± 7.3 | 33.7 ± 6.6 | 37.5 ± 15.5 | 9.7 ± 7.6 | 46.1 | 0.71 | 0.73 | 1.55 | 1.71 |
| CR1 | Lizard | 1 | 53.8 | 46.2 | 50.2 | 37.8 | 50.7 | 2.0 | 52.4 | 0.78 | 0.78 | 1.15 | 1.38 |
|  | Frog | 3 | 58.2 ± 1.0 | 41.8 ± 1.0 | 46.1 ± 1.3 | 36.0 ± 1.3 | 42.6 ± 0.6 | 5.0 ± 1.0 | 50.1 | 0.85 | 0.85 | 1.17 | 1.86 |
|  | Fish | 8 | 65.8 ± 3.5 | 34.2 ± 3.5 | 37.7 ± 2.6 | 31.1 ± 4.7 | 32.5 ± 3.5 | 8.5 ± 3.2 | 42.4 | 0.75 | 0.76 | 1.79 | 2.02 |
| L2 | Lizard | 17 | 45.1 ± 1.2 | 54.9 ± 1.2 | 59.1 ± 1.0 | 46.1 ± 0.8 | 59.4 ± 2.9 | 0.9 ± 0.7 | 54.7 | 0.77 | 0.78 | 1.11 | 1.37 |
|  | Frog | 2 | 52.0 ± 0.9 | 48.0 ± 0.9 | 49.7 ± 0.8 | 46.1 ± 0.2 | 48.4 ± 1.6 | 2.5 ± 0.7 | 49.9 | 0.79 | 0.81 | 1.31 | 1.44 |
|  | Fish | 19 | 57.7 ± 4.4 | 42.3 ± 4.4 | 45.6 ± 3.8 | 38.5 ± 4.0 | 40.4 ± 6.6 | 3.1 ± 2.7 | 51.7 | 0.74 | 0.77 | 1.31 | 1.49 |
| Rex1 | Frog | 3 | 51.1 ± 1.2 | 48.9 ± 1.2 | 48.7 ± 1.6 | 43.6 ± 0.5 | 55.0 ± 1.4 | 1.3 ± 0.6 | 54.6 | 0.84 | 0.83 | 1.089 | 1.927 |
|  | Fish | 12 | 47.7 ± 4.9 | 52.3 ± 4.9 | 47.6 ± 2.9 | 42.5 ± 2.8 | 47.7 ± 10.1 | 1.7 ± 2.8 | 52.6 | 0.77 | 0.79 | 1.19 | 1.48 |
| I | Lizard | 1 | 54.6 | 43.4 | 50.2 | 42.3 | 43.7 | 4 | 53.2 | 0.76 | 0.76 | 1.20 | 1.48 |
|  | Fish | 6 | 65.7 ± 1.5 | 34.3 ± 1.5 | 41.0 ± 1.6 | 33.4 ± 0.8 | 28.4 ± 2.7 | 15.8 ± 6.2 | 42.4 | 0.72 | 0.74 | 1.67 | 1.80 |
| R4 | Lizard | 1 | 57.3 | 43.7 | 47.5 | 34.3 | 46.4 | 2 | 51.1 | 0.79 | 0.77 | 1.17 | 1.44 |
|  | Fish | 1 | 47.5 | 52.5 | 51.3 | 43.2 | 63.1 | 1 | 49.5 | 0.70 | 0.73 | 1.27 | 1.40 |
| RTE | Lizard | 1 | 50.0 | 50.0 | 54.3 | 41.2 | 54.5 | 0 | 53.3 | 0.79 | 0.79 | 1.12 | 1.36 |
|  | Fish | 5 | 45.4 ± 6.1 | 54.6 ± 6.1 | 57.1 ± 5.3 | 46.4 ± 6.4 | 60.4 ± 7.3 | 0.4 ± 0.5 | 50.7 | 0.72 | 0.74 | 1.27 | 1.45 |
| Penelope | Lizard | 7 | 62.7 ± 3.4 | 37.3 ± 3.4 | 40.5 ± 4.0 | 34.8 ± 0.9 | 36.5 ± 5.7 | 5.4 ± 2.6 | 49 | 0.75 | 0.75 | 1.40 | 1.61 |
|  | Frog | 11 | 56.4 ± 2.9 | 43.4 ± 2.9 | 46.0 ± 2.0 | 39.7 ± 2.3 | 44.1 ± 5.0 | 1.5 ± 1.3 | 52.7 | 0.84 | 0.84 | 1.15 | 1.39 |
|  | Fish | 3 | 55.2 ± 1.5 | 44.8 ± 1.5 | 46.4 ± 3.7 | 37.1 ± 1.0 | 50.7 ± 0.2 | 5.0 ± 5.0 | 54.1 | 0.68 | 0.69 | 1.24 | 1.62 |

Elements of the L1, CR1, I and Penelope clades show a clear tendency to be AT-rich, although there are notable differences between nLTR-RT clades within the same genomes. For instance, in lizard, L1 is on average 67% AT while CR1 is 54% AT. The same clade can also differ substantially in base composition among hosts as demonstrated for CR1, whose nucleotide content ranges from 54% AT in lizard to 66% in fish. In L1, there is a strong bias in favor of A on the positive strand in lizard and mammals (43 and 42% A *versus* 24 and 21% T, respectively), a more moderate bias in the frog (33% A and 23% T) and no bias in fish, where A (33%) and T (31%) are represented almost equally. In CR1, there is an A bias in lizard but not in frog and fish. The base composition of Penelope and I elements always showed an A bias on the positive strand.

The Rex1, L2 and RTE show distinct patterns. The base composition of Rex1 is similar to the base composition of the exome of the source species and does not differ between frog and fish. The base composition of the RTE clade tends to be GC-rich in fish and lizard, but there are substantial differences among families with proportions of GC ranging from 48% to 61%. Interestingly, RTE families can differ considerably in nucleotide content within the same organism. This is exemplified in medaka, whose genome hosts 3 families of RTE. RTE 2 and 3 have GC content just below 50% (48 and 47%, respectively), while RTE1 contains 60% GC (supplementary material 1). All fish RTE elements show a G bias on the positive strand (33 to 39% G). The base composition of the L2 clade is equally disparate. L2 is GC-rich in lizard (55% GC), AT-rich in fish (58% AT) and moderately AT-rich in frog (51% AT), C (34%) and then T (30%) being the most represented bases in this organism. It should be noted that the base composition within each vertebrate lineage shows little variation as indicated by small standard deviations on figure 2 and table 1.

We extended our analysis to dinucleotides (table 2) since genomes exhibit variations in their usage of dinucleotides [34]. We present here the frequencies of dinucleotides in three clades, L1, CR1 and L2, since these clades reflect the range of base composition in nLTR-RT. In general, the frequency of the different dinucleotides reflects the base composition of the elements. Elements that are AT-rich, such as L1, CR1 and the fish L2, are also enriched in the dinucleotides ApA, ApT, TpA and TpT, while there is a paucity of GC-rich dinucleotides. In the lizard L2, the most abundant dinucleotides are expectedly GC-rich, the four most represented dinucleotides being CpC, CpT, GpG and TpG. The frog L2 is somewhat unusual: the CpT and TpC are abundant, which is consistent with the base composition of the elements, but the next two most common dinucleotides are surprisingly ApG and GpA, although A and G represent respectively only 22% and 14% of nucleotides. We then assessed if the frequency of each dinucleotide deviates from the expected frequency given the abundance of the nucleotides in the sequence (table 2). Overall the observed and expected frequencies are remarkably close (as demonstrated by the ratios between observed and expected dinucleotide frequencies), demonstrating that the base composition dictates the frequency of dinucleotides. There are however three dinucleotides that are systematically under-represented across species and across clades, CpG, GpT and TpA, suggesting universal selection against these dinucleotides. The only clade that shows substantial deviation from expectation is L2 in frog. Here ApC, CpC, CpT, TpC and TpT are substantially under-represented (although C and T are the most abundant nucleotides) but ApG, GpA and GpG are over-represented given the frequency of the constitutive nucleotides.

**Table 2.**
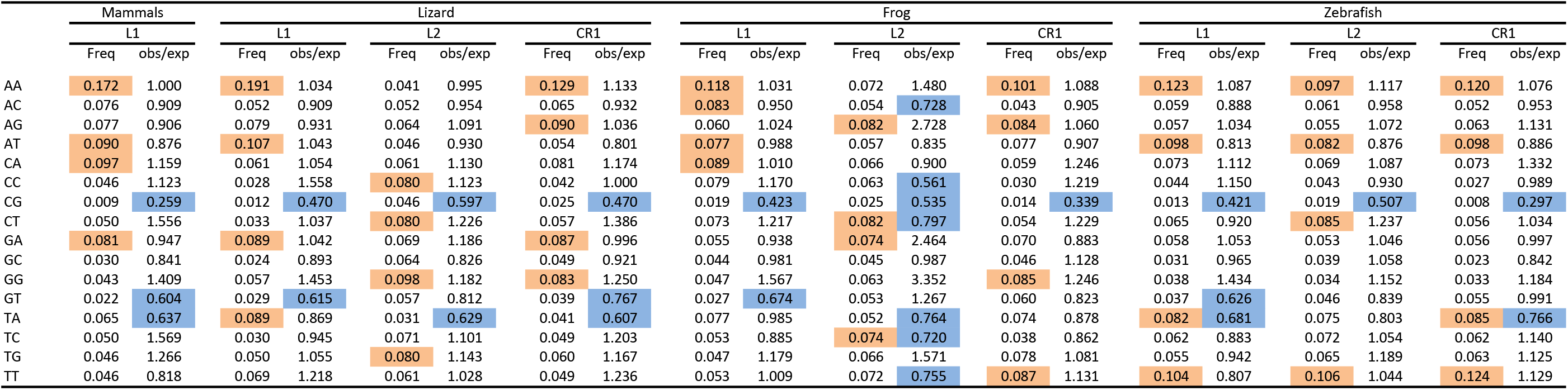
Frequency of nucleotides and ratio between the observed and expected frequencies. Four each clade the four most common dinucleotides are highlighted in orange. Ratios that are lower than 0.80 are highlighted in blue.

### Differences in base composition reflect long-term evolutionary trends

To interpret long-term evolutionary changes in base composition and the possible impact of horizontal gene transfer (HGT), we investigated differences in base composition in a phylogenetic context. To this end, we built phylogenetic trees for the major clades of nLTR-RT (figures 3 to 6) and we estimated the level of identity among amino acid sequences, within and between clade.

**Figure 3.**
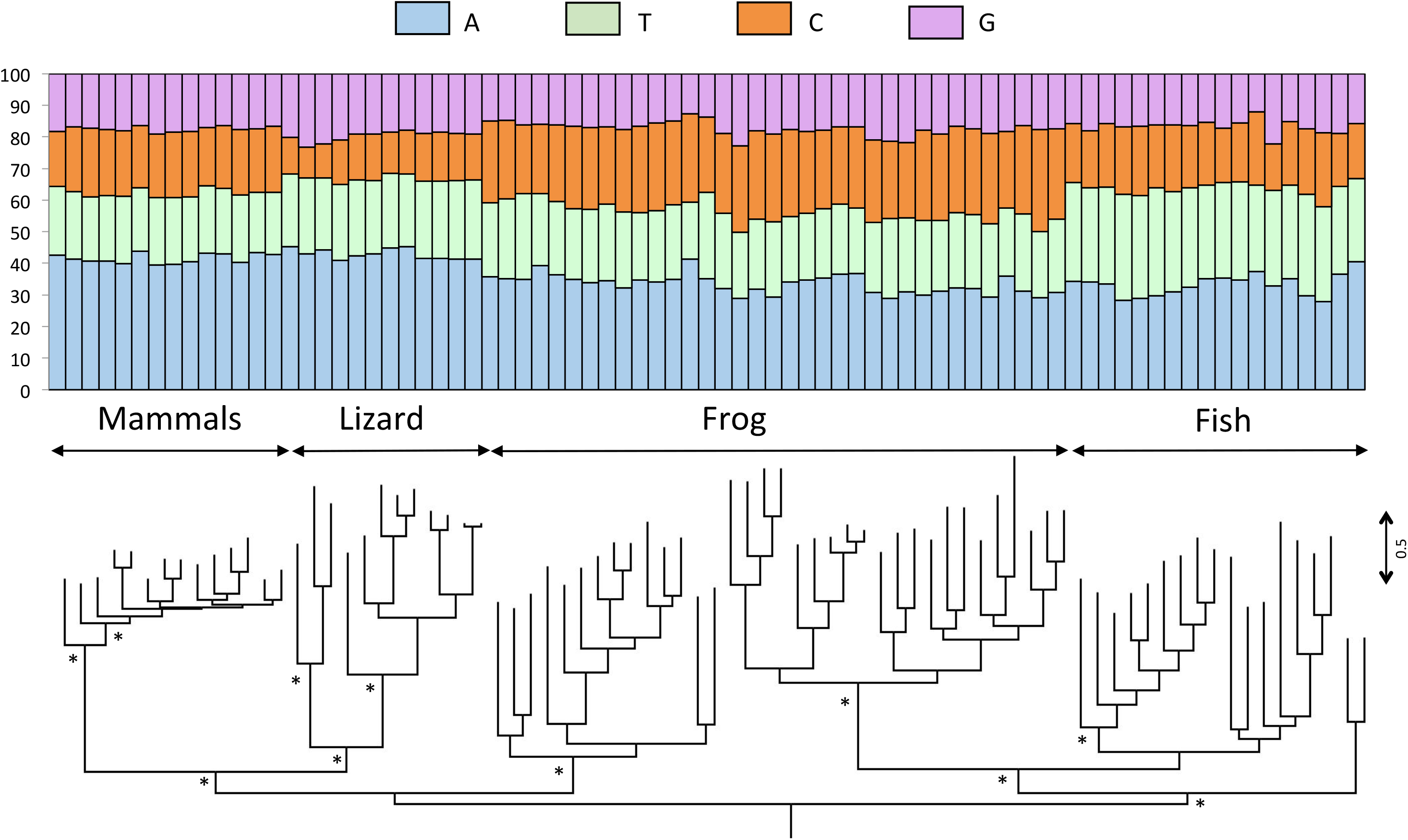
Phylogenetic relationships among L1 elements based on the entire ORF2. The base composition of each element is shown above the tree. The tree was built with the maximum likelihood method using the LG+G+I+F model of mutation and its robustness was assessed by 500 bootstrap replicates. Nodes that are supported by bootstrap values higher than 70% are indicated with an asterisk.

The L1 clade consists of 5 sub-clades: a lizard clade, a mammalian clade, a fish clade (which includes all fish sequences but two) and two frog clades that are not sister to each other (figure 3). The topology of the tree suggests the persistence in frog and in fish of ancient and diverse L1 lineages, whose divergence predates the split between teleost fish and tetrapods. The amino acid identity among the most divergent L1 sequences within the lizard, fish and frog clades are low, with average values of 26.5, 27.5 and 31.2% respectively, suggesting that multiple lineages of L1 can coexist and evolve for extended periods of evolutionary time within the same genome, as previously reported in [16, 35, 36]. Despite this high level of divergence, the base composition remains remarkably constant within vertebrate lineage (figure 3), which is consistent with the small standard deviations on figure 2. This suggests some long-term selective pressure or functional constraint on L1 to maintain AT-richness (in all vertebrates) and an A bias on the positive strand (in mammals, lizard and frog).

Elements classified as CR1 or L2 form two major clades and each contain additional sub-clades (figure 4). The CR1 clade consists of three fish lineages (which do not form a clade) and a tetrapod CR1 cluster (figure 4). Tetrapod CR1 and most fish CR1 elements have an ORF upstream of ORF2. In tetrapod this ORF contains an esterase domain while we failed to identify any known conserved motif or domain in fish. The base composition is homogenous within each of these sub-clades, with a moderate AT-bias in tetrapods and a strong bias in all fish (zebrafish, medaka, stickleback). The L2 clade is more complex and consists of six sub-clades (numbered I through VI on figure 4): a lizard sub-clade (I), a fish/frog sub-clade (II) and 4 fish sub-clades (III to VI). Elements belonging to the fish/frog sub-clade and to two of the fish sub-clades (sub-clades II to IV) have a single ORF (ORF2) while the other two fish sub-clades (V and VI) have acquired an additional ORF containing an esterase domain. Since these two di-cistronic sub-clades are not sister to each other, it is possible that they have acquired a second ORF independently. The two di-cistronic sub-clades V and VI harbor the strong AT bias observed for other fish elements (~61% AT) while elements in the mono-cistronic sub-clades (II and III) contain a large proportion of the nucleotides C and T and elements in clade IV have a T bias, the three other bases being equally represented. Each of the fish sub-clades contains sequences from multiple species and the level of identity between species in each sub-clade is similar (~50% identity). This suggests that the base composition has been maintained in those genomes since before the species diverged. The lizard L2 sub-clades consist of elements that are all GC-rich. The lizard L2 clade experienced an intense diversification and a number of closely related families are concurrently active in this genome [36]. However, the most divergent subgroups in this sub-clade are on average 36.6% identical at the amino acid level indicating that the GC-richness of L2 in lizard is ancient and has persisted through extended periods of evolutionary time. Most, but not all, lizard L2s have a second ORF, for which we failed to find any conserved domain. It appears that the lizard L2 acquired a second ORF independently twice since the ORF of the most basal elements does not share any homology with the ORF found in the families that experienced a recent diversification.

**Figure 4.**
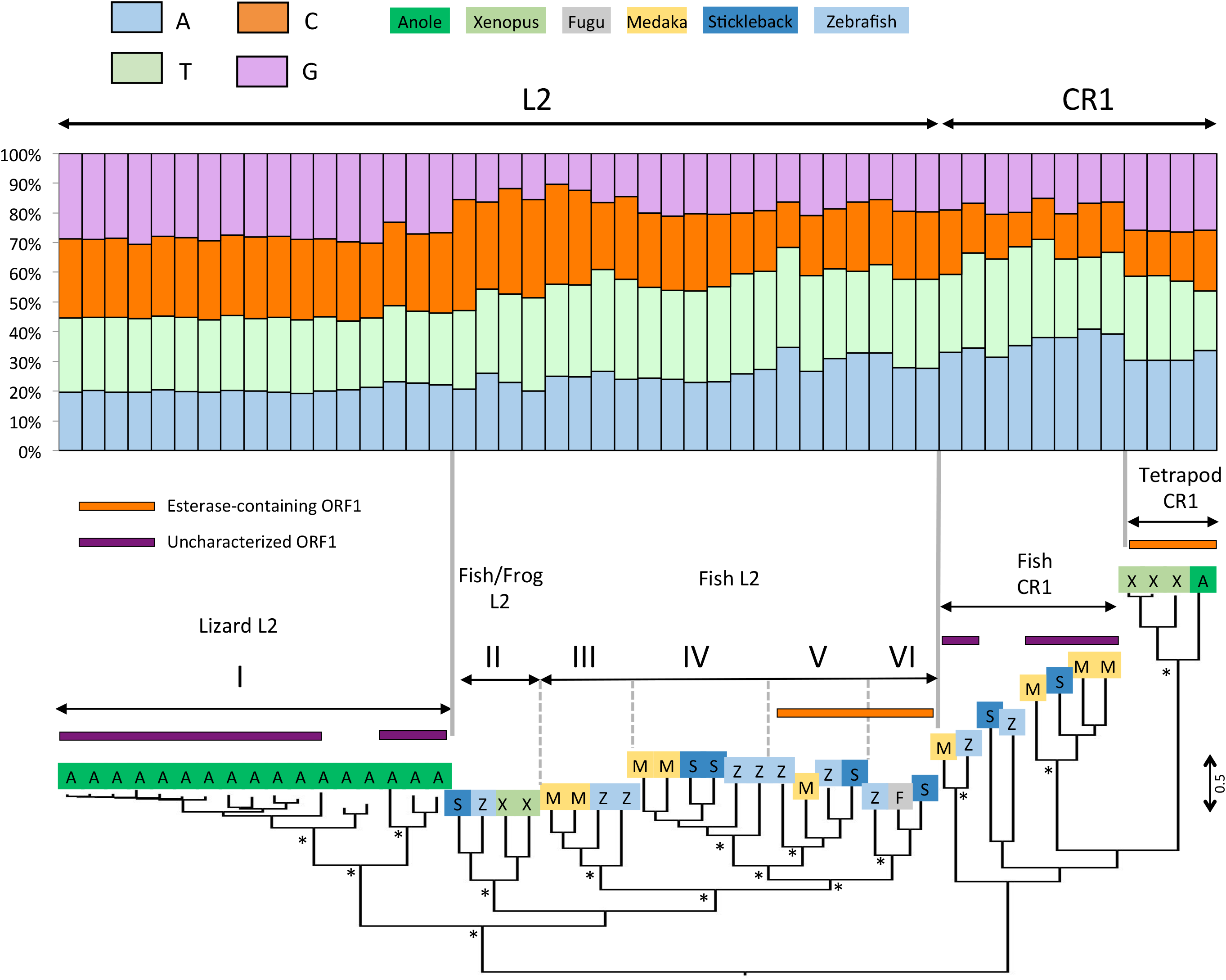
Phylogenetic relationships among CR1 and L2 elements based on the entire ORF2. The base composition of each element is shown above the tree. The tree was built with the maximum likelihood method using the LG+G+I+F model of mutation and its robustness was assessed by 500 bootstrap replicates. Nodes that are supported by bootstrap values higher than 70% are indicated with an asterisk. Roman numbers I to VI refer to the L2 sub-clades (see text).

The Tx1 clade consists of three highly divergent sub-clades (named I through III on figure 5). Each of these three sub-clades remains monophyletic in a wider phylogenetic context (data not shown). In all three sub-clades AT-rich elements dominate but four nested sequences, three in sub-clade I and one in sub-clade III (indicated with arrows on figure 5), are GC rich (~53%). These sequences do not group together and are derived from three different hosts (fugu, stickleback and frog), while all other sequences but one come from zebrafish. This pattern suggests that host-specific forces can cause changes in base composition, or that these four sequences were transferred horizontally from an organism that harbors Tx1 elements with different base composition.

**Figure 5.**
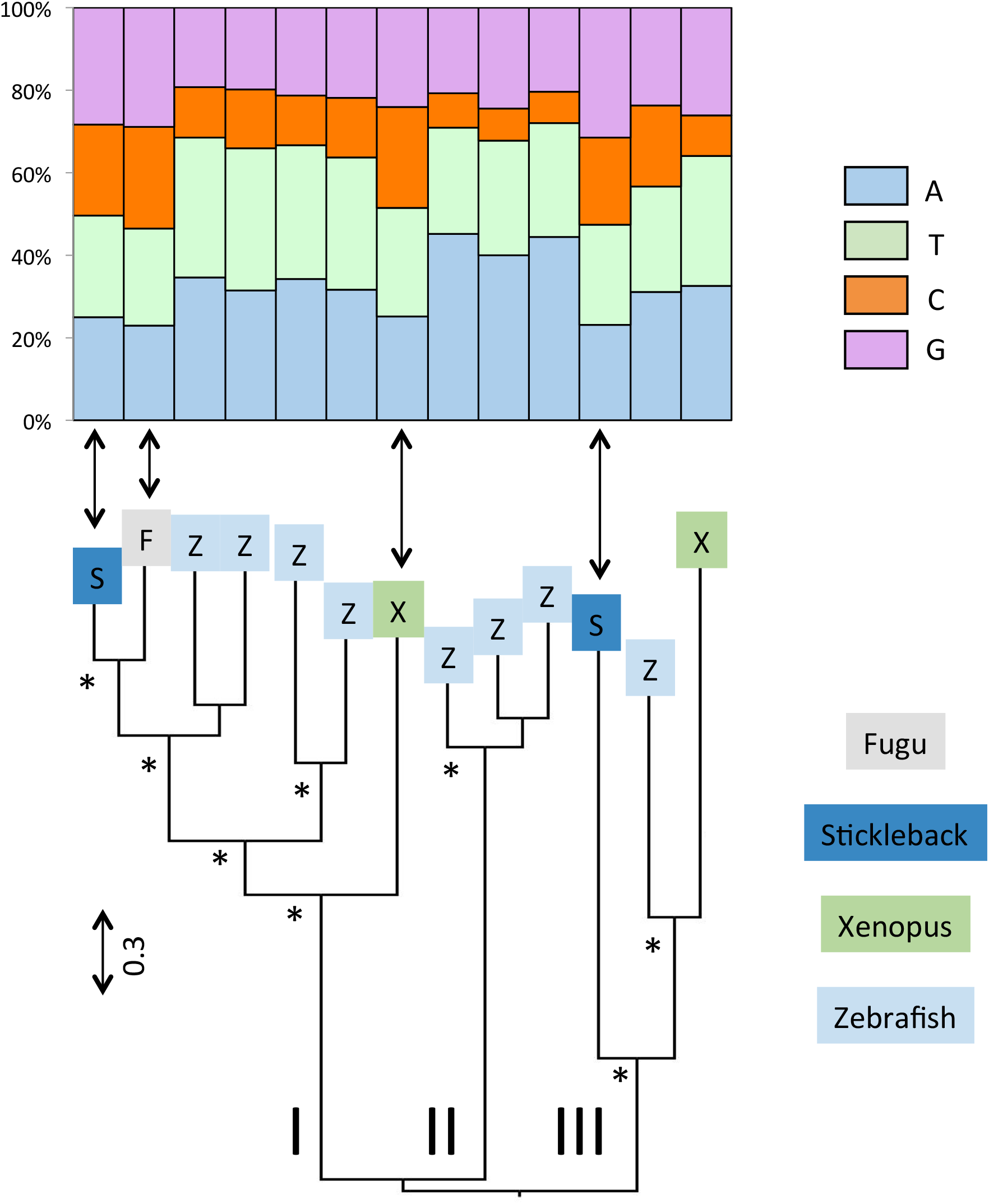
Phylogenetic relationships among Tx1 elements based on the entire ORF2. The base composition of each element is shown above the tree. The tree was built with the maximum likelihood method using the LG+G+I+F model of mutation and its robustness was assessed by 500 bootstrap replicates. Nodes that are supported by bootstrap values higher than 70% are indicated with an asterisk. Elements with comparatively low AT-content are indicated with arrows (see text).

In contrast to the clades described above, the AT-rich base composition of Rex1, I and Penelope has been conserved over extended periods of evolutionary time. For instance, the diversification of Rex1 predates the split between fish and tetrapods and it has persisted in both fish and frog (figure 6). Yet, despite this ancient history, the base composition of Rex1 has remained constant over a timespan of 525My [37]. It should be noted however that one of the frog sequences (indicated with an asterisk on figure 6) is much more similar to its closest fish sequence (78% identity) than expected, which is suggestive of an ancient event of HGT between frogs and fish, or in both lineages from a common source. Thus, we can’t exclude that the HGT of Rex1 among lineages has contributed to the apparent pattern of homogeneity in base composition among organisms, although the divergences between elements in the rest of the tree are consistent with the host divergence and suggest that vertical transfer is the main mode of transmission of Rex1.

**Figure 6.**
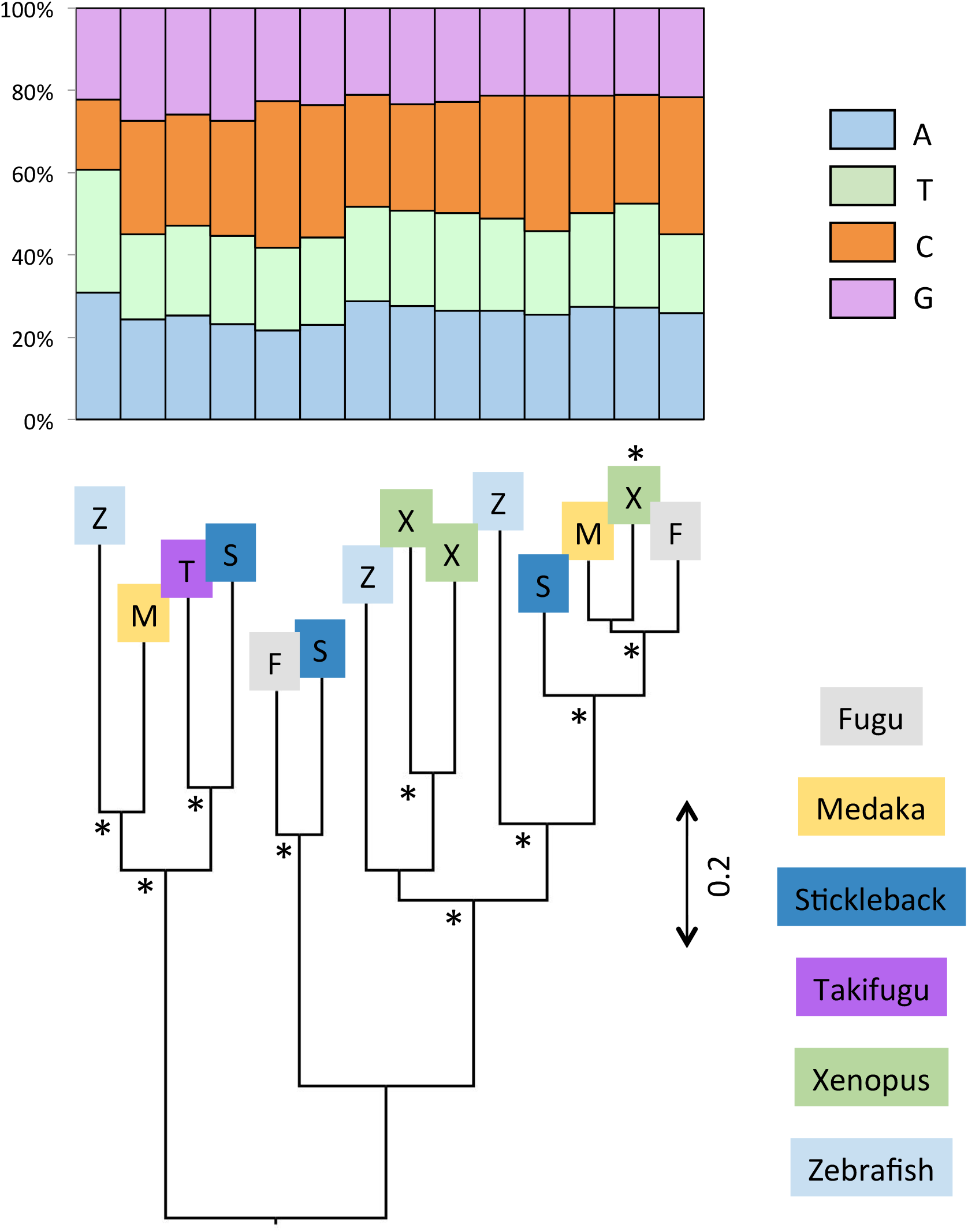
Phylogenetic relationships among Rex1 elements based on the entire ORF2. The base composition of each element is shown above the tree. The tree was built with the maximum likelihood method using the LG+G+I+F model of mutation and its robustness was assessed by 500 bootstrap replicates. Nodes that are supported by bootstrap values higher than 70% are indicated with an asterisk. The frog element suspected of HGT is indicated with an asterisk.

### Variation in base composition across the ORFs

In order to determine if the difference in base composition extends outside the ORF containing the reverse transcriptase domain (ORF2), we compared the base composition of ORF2 with the upstream ORFs (ORF1) in those clades that have two ORFs, such as L1, most CR1 and some L2 elements (figure 7). For L1 and CR1, ORF1 shows the same nucleotide bias as ORF2 but the bias always tends to be stronger in ORF2 than in ORF1, i.e. ORF2 is always richer in AT than ORF1, and this is true in all vertebrates. In contrast, the two ORFs of L2 differ markedly in nucleotide content. In lizard, ORF2 is GC-rich (~55% GC) but ORF1 is AT-rich (54%). In fish, the phylogenetic analysis suggests that ORF1 was independently acquired twice since the two sub-clades containing elements with an ORF1 are not sister to each other. This is supported by the fact that the base composition of their ORFs differs, 55% AT in one of the sub-clades and 45% in the other one, although their ORF2 is equally AT-rich (61% AT). It should be noted that since ORF1 is not homologous among CR1 and L2 sub-clades, variations in base composition may not result from long term processes acting on base composition but instead may reflect the original nucleotide content of the sequence that was recruited by the element to form a novel ORF.

**Figure 7.**
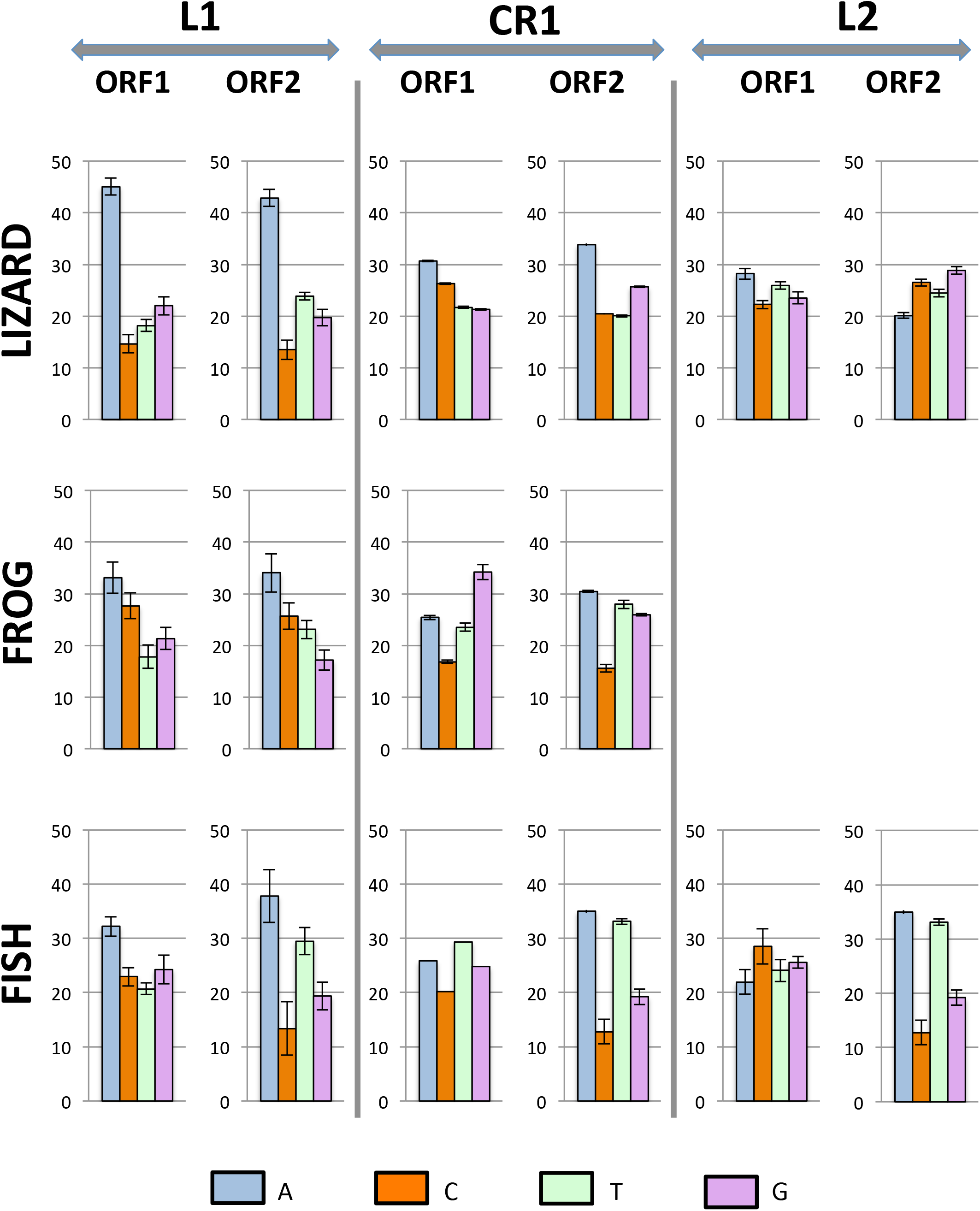
Comparison of the nucleotide content of ORF1 and ORF2 in L1, CR1 and L2.

### Variation in base composition at the codon level

We then investigated base composition at the codon-level in ORF2 (figure 8). The first two positions of codons determine the encoded amino acid but the base composition at the third position, which is mostly neutral, should reflect the mutational process alone. In L1, the base composition is similar at the three position of codons, showing a strong A bias in mammals and lizard, a moderate one in frog and equal contribution of A and T in fish. In CR1, the first and second positions of the codons show similar composition among all organisms. The two most abundant bases are A and then G at the first position, and A and T at the second position, demonstrating long-term selective constrains with regard to the encoded amino acid. Composition at the mostly neutral third position varies, with similar frequency of AT and GC in lizard, a moderate AT bias in frog (with T the most abundant base) and a strong AT bias in fish. In L2, the difference between species is notable. In frog, the three positions are enriched in C and T. In lizard the first and third positions are GC-rich (59% at both positions) while T is the most common base at the second position, followed by C. In fish, L2 tends to be AT-rich at the three positions but there is an A bias at the first position (31% A), a T bias at the second (33%) and third (36%) position. It is interesting to note that the base composition at the third, mostly neutral, position differs among clades between species and among clades within the same host. This suggests that a strictly neutral process does not drive the base composition at the third position and that selection is acting on base composition, independently of the protein-coding capability of the sequence.

**Figure 8.**
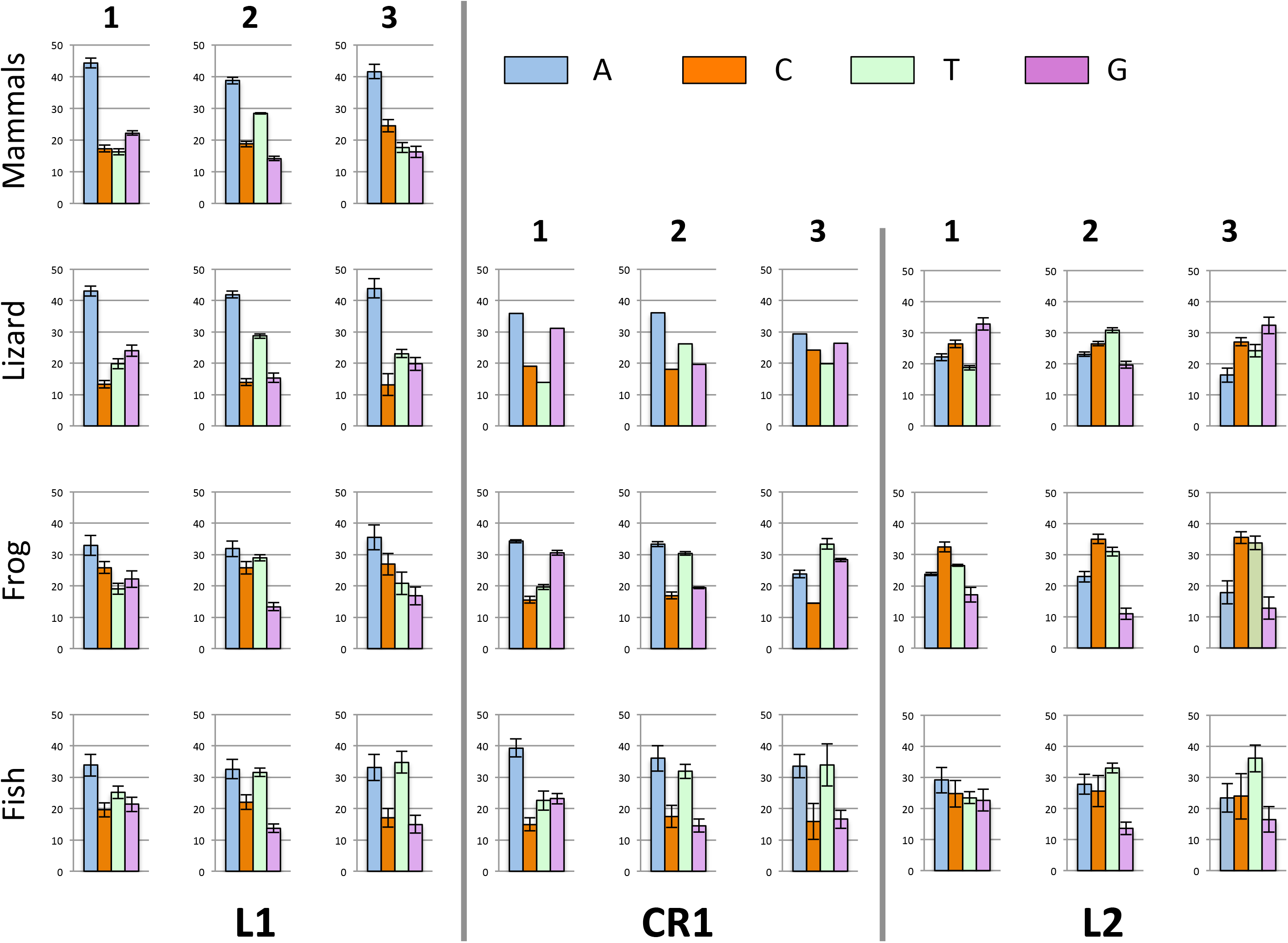
Base composition at the three codon positions in L1, CR1 and L2 ORF2.

We then investigated how the base composition at the three positions affects codon usage and the amino acid sequence of the encoded protein. Figure 9 shows examples of codon usage for L1 and L2 ORF2 in lizard and zebrafish, compared with the codon usage of the hosts’ exomes. For lizard L1, it is always the A-rich codon that is preferred while the opposite trend is found for L2. For instance, codon GAA is used more than 70% of the time in L1 to encode glutamic acid (E on figure 9), whereas it is GAG that is used more than 80% of the time in L2, and both codons are used almost equally in the host exome. In contrast, there is very little difference in codon usage between L1 and L2 in fish since both elements are similarly AT-rich and both L1 and L2 show a preference for AT-rich codons compared with the exome. These trends are summarized by the Relative Synonymous Codon Usage (RSCU; table 3), which shows a preference for codons with an A at the third position in AT-rich elements with an A bias on the positive strand and a preference for an A or a T for the fish AT-rich elements.

**Figure 9.**
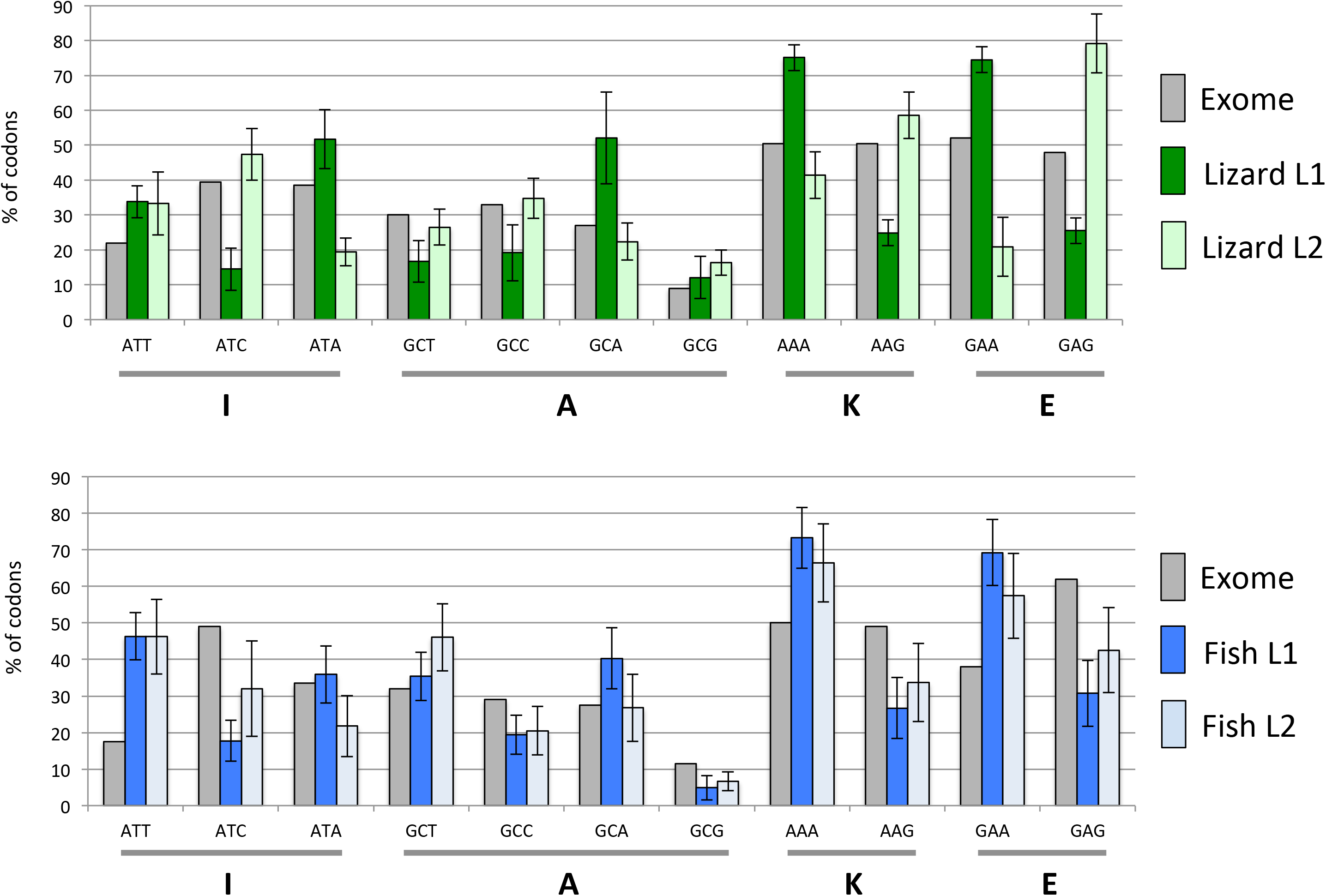
Codon usage in ORF2 for four amino acids in fish and lizard L1 and L2.

**Table 3.**
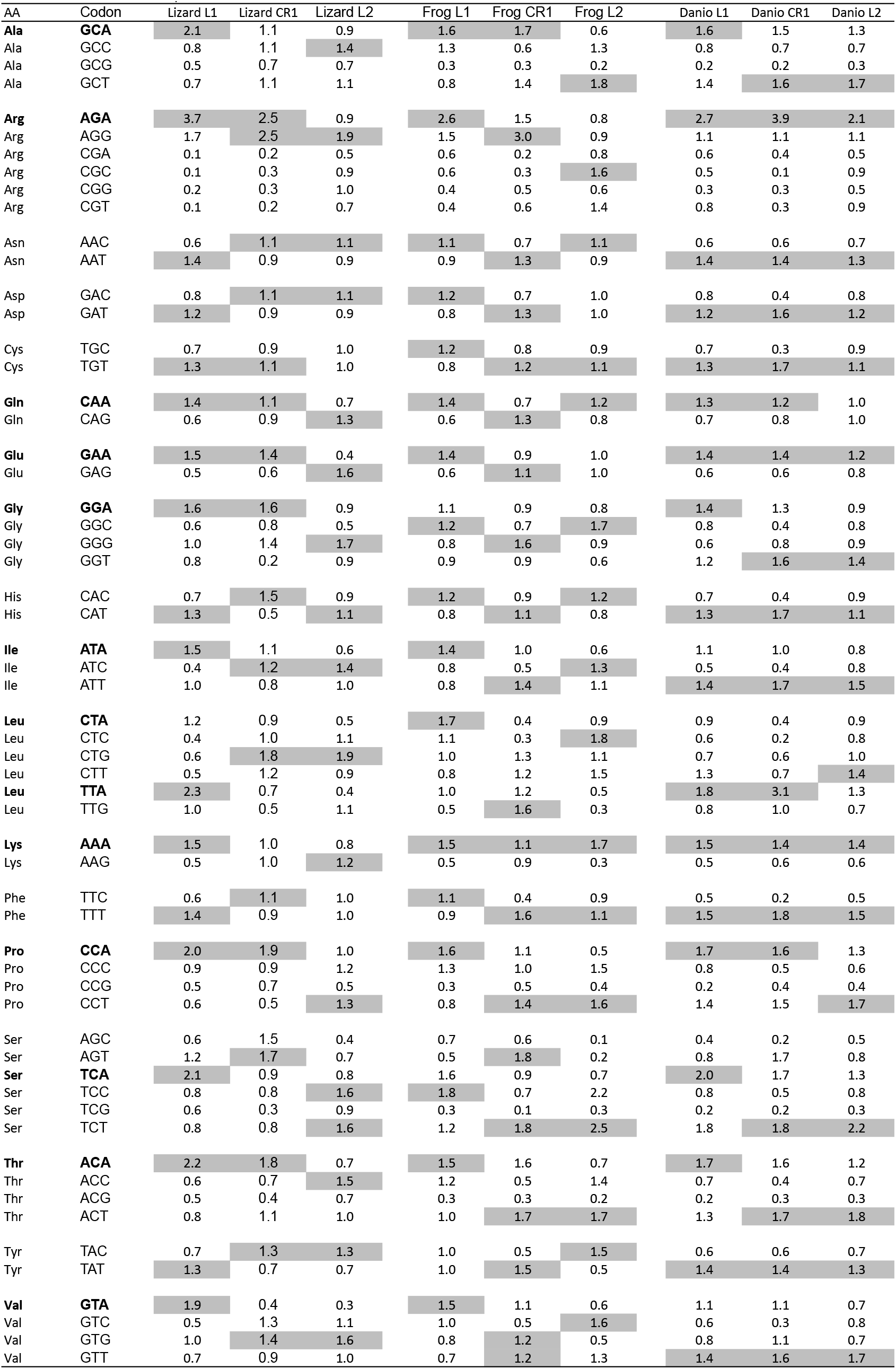
Relative Codon Usage (RSCU) for ORF2 in lizard, frog and fish L1, CR1 and L2 elements. The highest RSCU for each amino acid is highlighted in grey. Codons with an A at the third position are in bold.

| AA | Codon | Lizard L1 | Lizard CR1 | Lizard L2 | Frog L1 | Frog CR1 | Frog L2 | Danio L1 | Danio CR1 | Danio L2 |
| --- | --- | --- | --- | --- | --- | --- | --- | --- | --- | --- |
| **Ala** | **GCA** | 2.1 | 1.1 | 0.9 | 1.6 | 1.7 | 0.6 | 1.6 | 1.5 | 1.3 |
| Ala | GCC | 0.8 | 1.1 | 1.4 | 1.3 | 0.6 | 1.3 | 0.8 | 0.7 | 0.7 |
| Ala | GCG | 0.5 | 0.7 | 0.7 | 0.3 | 0.3 | 0.2 | 0.2 | 0.2 | 0.3 |
| Ala | GCT | 0.7 | 1.1 | 1.1 | 0.8 | 1.4 | 1.8 | 1.4 | 1.6 | 1.7 |
| **Arg** | **AGA** | 3.7 | 2.5 | 0.9 | 2.6 | 1.5 | 0.8 | 2.7 | 3.9 | 2.1 |
| Arg | AGG | 1.7 | 2.5 | 1.9 | 1.5 | 3.0 | 0.9 | 1.1 | 1.1 | 1.1 |
| Arg | CGA | 0.1 | 0.2 | 0.5 | 0.6 | 0.2 | 0.8 | 0.6 | 0.4 | 0.5 |
| Arg | CGC | 0.1 | 0.3 | 0.9 | 0.6 | 0.3 | 1.6 | 0.5 | 0.1 | 0.9 |
| Arg | CGG | 0.2 | 0.3 | 1.0 | 0.4 | 0.5 | 0.6 | 0.3 | 0.3 | 0.5 |
| Arg | CGT | 0.1 | 0.2 | 0.7 | 0.4 | 0.6 | 1.4 | 0.8 | 0.3 | 0.9 |
| Asn | AAC | 0.6 | 1.1 | 1.1 | 1.1 | 0.7 | 1.1 | 0.6 | 0.6 | 0.7 |
| Asn | AAT | 1.4 | 0.9 | 0.9 | 0.9 | 1.3 | 0.9 | 1.4 | 1.4 | 1.3 |
| Asp | GAC | 0.8 | 1.1 | 1.1 | 1.2 | 0.7 | 1.0 | 0.8 | 0.4 | 0.8 |
| Asp | GAT | 1.2 | 0.9 | 0.9 | 0.8 | 1.3 | 1.0 | 1.2 | 1.6 | 1.2 |
| Cys | TGC | 0.7 | 0.9 | 1.0 | 1.2 | 0.8 | 0.9 | 0.7 | 0.3 | 0.9 |
| Cys | TGT | 1.3 | 1.1 | 1.0 | 0.8 | 1.2 | 1.1 | 1.3 | 1.7 | 1.1 |
| **Gln** | **CAA** | 1.4 | 1.1 | 0.7 | 1.4 | 0.7 | 1.2 | 1.3 | 1.2 | 1.0 |
| Gln | CAG | 0.6 | 0.9 | 1.3 | 0.6 | 1.3 | 0.8 | 0.7 | 0.8 | 1.0 |
| **Glu** | **GAA** | 1.5 | 1.4 | 0.4 | 1.4 | 0.9 | 1.0 | 1.4 | 1.4 | 1.2 |
| Glu | GAG | 0.5 | 0.6 | 1.6 | 0.6 | 1.1 | 1.0 | 0.6 | 0.6 | 0.8 |
| **Gly** | **GGA** | 1.6 | 1.6 | 0.9 | 1.1 | 0.9 | 0.8 | 1.4 | 1.3 | 0.9 |
| Gly | GGC | 0.6 | 0.8 | 0.5 | 1.2 | 0.7 | 1.7 | 0.8 | 0.4 | 0.8 |
| Gly | GGG | 1.0 | 1.4 | 1.7 | 0.8 | 1.6 | 0.9 | 0.6 | 0.8 | 0.9 |
| Gly | GGT | 0.8 | 0.2 | 0.9 | 0.9 | 0.9 | 0.6 | 1.2 | 1.6 | 1.4 |
| His | CAC | 0.7 | 1.5 | 0.9 | 1.2 | 0.9 | 1.2 | 0.7 | 0.4 | 0.9 |
| His | CAT | 1.3 | 0.5 | 1.1 | 0.8 | 1.1 | 0.8 | 1.3 | 1.7 | 1.1 |
| **Ile** | **ATA** | 1.5 | 1.1 | 0.6 | 1.4 | 1.0 | 0.6 | 1.1 | 1.0 | 0.8 |
| Ile | ATC | 0.4 | 1.2 | 1.4 | 0.8 | 0.5 | 1.3 | 0.5 | 0.4 | 0.8 |
| Ile | ATT | 1.0 | 0.8 | 1.0 | 0.8 | 1.4 | 1.1 | 1.4 | 1.7 | 1.5 |
| **Leu** | **CTA** | 1.2 | 0.9 | 0.5 | 1.7 | 0.4 | 0.9 | 0.9 | 0.4 | 0.9 |
| Leu | CTC | 0.4 | 1.0 | 1.1 | 1.1 | 0.3 | 1.8 | 0.6 | 0.2 | 0.8 |
| Leu | CTG | 0.6 | 1.8 | 1.9 | 1.0 | 1.3 | 1.1 | 0.7 | 0.6 | 1.0 |
| Leu | CTT | 0.5 | 1.2 | 0.9 | 0.8 | 1.2 | 1.5 | 1.3 | 0.7 | 1.4 |
| **Leu** | **TTA** | 2.3 | 0.7 | 0.4 | 1.0 | 1.2 | 0.5 | 1.8 | 3.1 | 1.3 |
| Leu | TTG | 1.0 | 0.5 | 1.1 | 0.5 | 1.6 | 0.3 | 0.8 | 1.0 | 0.7 |
| **Lys** | **AAA** | 1.5 | 1.0 | 0.8 | 1.5 | 1.1 | 1.7 | 1.5 | 1.4 | 1.4 |
| Lys | AAG | 0.5 | 1.0 | 1.2 | 0.5 | 0.9 | 0.3 | 0.5 | 0.6 | 0.6 |
| Phe | TTC | 0.6 | 1.1 | 1.0 | 1.1 | 0.4 | 0.9 | 0.5 | 0.2 | 0.5 |
| Phe | TTT | 1.4 | 0.9 | 1.0 | 0.9 | 1.6 | 1.1 | 1.5 | 1.8 | 1.5 |
| **Pro** | **CCA** | 2.0 | 1.9 | 1.0 | 1.6 | 1.1 | 0.5 | 1.7 | 1.6 | 1.3 |
| Pro | CCC | 0.9 | 0.9 | 1.2 | 1.3 | 1.0 | 1.5 | 0.8 | 0.5 | 0.6 |
| Pro | CCG | 0.5 | 0.7 | 0.5 | 0.3 | 0.5 | 0.4 | 0.2 | 0.4 | 0.4 |
| Pro | CCT | 0.6 | 0.5 | 1.3 | 0.8 | 1.4 | 1.6 | 1.4 | 1.5 | 1.7 |
| Ser | AGC | 0.6 | 1.5 | 0.4 | 0.7 | 0.6 | 0.1 | 0.4 | 0.2 | 0.5 |
| Ser | AGT | 1.2 | 1.7 | 0.7 | 0.5 | 1.8 | 0.2 | 0.8 | 1.7 | 0.8 |
| **Ser** | **TCA** | 2.1 | 0.9 | 0.8 | 1.6 | 0.9 | 0.7 | 2.0 | 1.7 | 1.3 |
| Ser | TCC | 0.8 | 0.8 | 1.6 | 1.8 | 0.7 | 2.2 | 0.8 | 0.5 | 0.8 |
| Ser | TCG | 0.6 | 0.3 | 0.9 | 0.3 | 0.1 | 0.3 | 0.2 | 0.2 | 0.3 |
| Ser | TCT | 0.8 | 0.8 | 1.6 | 1.2 | 1.8 | 2.5 | 1.8 | 1.8 | 2.2 |
| **Thr** | **ACA** | 2.2 | 1.8 | 0.7 | 1.5 | 1.6 | 0.7 | 1.7 | 1.6 | 1.2 |
| Thr | ACC | 0.6 | 0.7 | 1.5 | 1.2 | 0.5 | 1.4 | 0.7 | 0.4 | 0.7 |
| Thr | ACG | 0.5 | 0.4 | 0.7 | 0.3 | 0.3 | 0.2 | 0.2 | 0.3 | 0.3 |
| Thr | ACT | 0.8 | 1.1 | 1.0 | 1.0 | 1.7 | 1.7 | 1.3 | 1.7 | 1.8 |
| Tyr | TAC | 0.7 | 1.3 | 1.3 | 1.0 | 0.5 | 1.5 | 0.6 | 0.6 | 0.7 |
| Tyr | TAT | 1.3 | 0.7 | 0.7 | 1.0 | 1.5 | 0.5 | 1.4 | 1.4 | 1.3 |
| **Val** | **GTA** | 1.9 | 0.4 | 0.3 | 1.5 | 1.1 | 0.6 | 1.1 | 1.1 | 0.7 |
| Val | GTC | 0.5 | 1.3 | 1.1 | 1.0 | 0.5 | 1.6 | 0.6 | 0.3 | 0.8 |
| Val | GTG | 1.0 | 1.4 | 1.6 | 0.8 | 1.2 | 0.5 | 0.8 | 1.1 | 0.7 |
| Val | GTT | 0.7 | 0.9 | 1.0 | 0.7 | 1.2 | 1.3 | 1.4 | 1.6 | 1.7 |

The codon usage bias was further investigated using three statistics - Nc, CAI and RCDI. Nc, also called the effective codon usage, which range from 20, when a single codon is used to encode for each amino acid, to 61 when all codons are used equally. We found the lowest values of Nc for elements that have the strongest A bias (mammalian or lizard L1) or AT bias (fish L1 and CR1) whereas elements that are GC rich (lizard L2) or have a base composition similar to the exome (Rex1) exhibit a higher value (table 1). This general trend is reflected by the fact that when the GC content at the third position of codon increases, so does the value of Nc (supplementary material 2).

The Codon Adaptation Index (CAI) is a measure of how closely the synonymous codon usage of a sequence matches that of a reference set, in our case the genome of the host. Table 1 shows the observed and expected average values of CAI given the base composition of the sequence. The values of CAI for the different types of nLTR-RT are remarkably similar to each other and the observed and expected values are almost identical. This indicates that there is no synonymous bias and that the frequency of the different codons fits what is expected given the nucleotide content of the sequence. This is consistent with the lack of correlation between the synonymous GC content and the observed values of CAI (supplementary material 2).

The Relative Codon Deoptimization Index (RCDI) is a measure of how different the codon usage in a sequence is relative to a reference set. An RCDI value of 1 indicates that the codon usage of a sequence is identical to the reference and the larger the value of RCDI the larger the difference in codon usage is. The lowest values of RCDI were found for elements with high GC content such as Rex1, the lizard L2, and RTE, and the highest values were found for the high AT rich elements such as L1 and the fish CR1 (table 1). In this case, we found a correlation between the GC content and the RCDI (supplementary material 2), which indicates that the higher the GC content the smaller the difference in the codon usage of the element and the codon usage of the host. Interestingly however, the observed RCDI is always lower than the expected RCDI, given the base composition of the sequence (table 1). This suggests that the codon usage of the element is closer to the codon usage of the host than expected given the base composition of its sequence, which indicates a certain level of codon usage adaptation.

Another consequence of nucleotide bias is that it can affect the amino acid composition of the ORFs, which in turn can affect the physico-chemical properties of the proteins as well as their stability. Figure 10 compares the amino acid composition of L1 and L2 ORF2 in lizard, frog and fish. The A-rich lizard L1 is considerably enriched in amino acids encoded by A-rich codons such as lysine (AAA and AAG) and Isoleucine (ATA, ATT and ATC), which respectively account for 13.6 and 9.8% of ORF2p. In contrast, amino acids encoded by GC rich codons, such as alanine (GCN; 8.6%) and arginine (CGN, AGT, AGC; 8.2%) are more abundant in lizard L2. Similarly, the CT-rich frog L2 encodes a protein enriched in CT-rich encoded amino acids, such as serine (TCN, AGT, AGC; 14.5%) and leucine (CTN, TTA, TTG; 15.9%). As expected, the amino acid composition of L1 and L2 in fish is very similar since these elements have similar nucleotide content.

**Figure 10.**
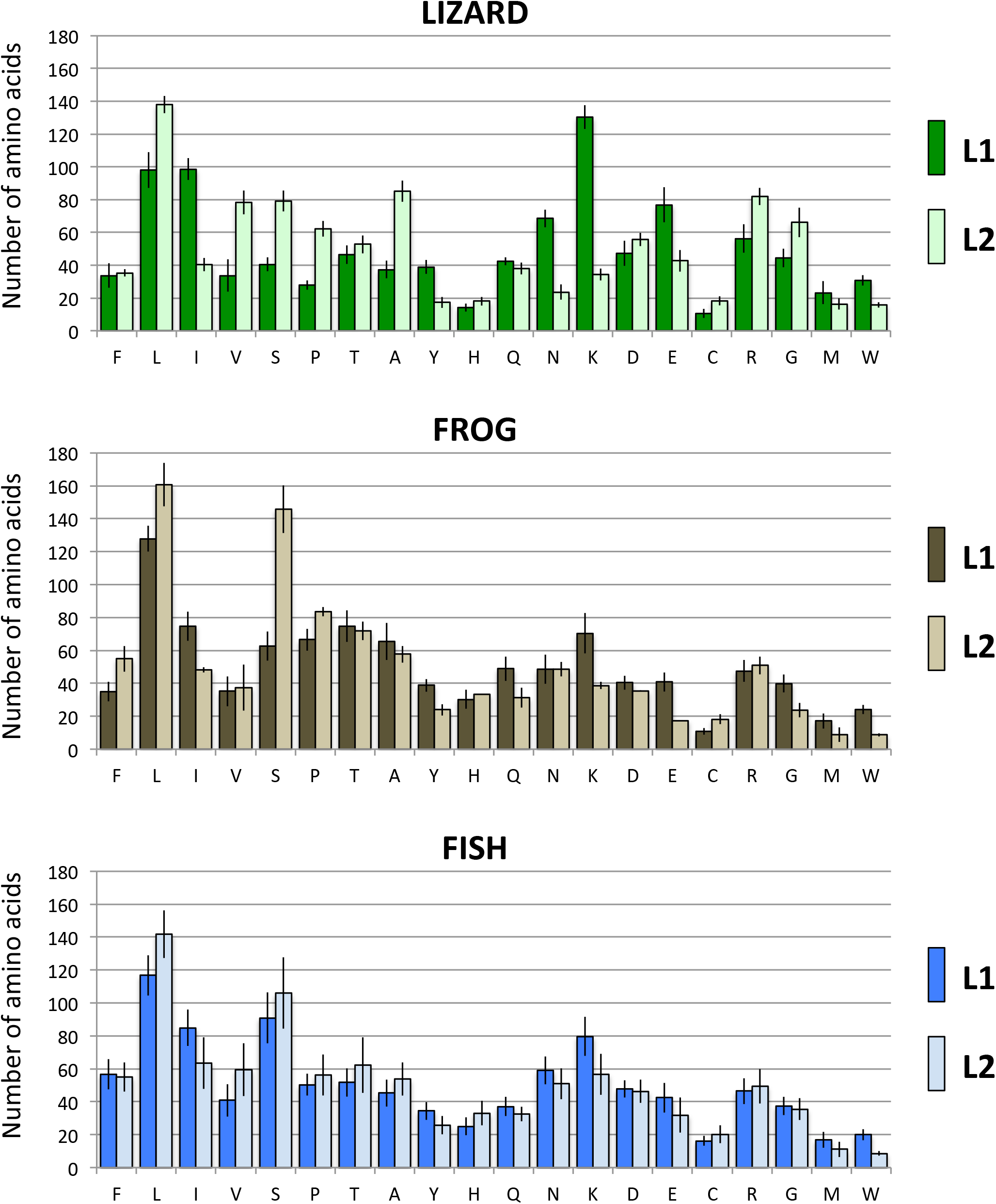
Number of amino acid in L1 and L2 ORF2 in lizard, frog and fish.

### Impact of base composition differences on the transcription

We then examined how biases in base composition can affect the transcription of retrotransposons. Because an AT-rich sequence is more likely to contain premature polyadenylation (polyA) signals, which would result in inefficient transcription [21], we assessed the number of canonical and non-canonical (AATAAA, ATTAAA) polyA signals in ORF2 (table 1). The number of polyA signals is correlated with the abundance in AT (supplementary material 3). AT rich elements, such as L1 have more polyA signals (up to 23 for ORF2 in lizard) than GC rich elements, which can have zero or one (such as RTE elements). This suggests that the ORFs of some clades may be transcribed much more efficiently than others. It should be noted that the number of polyA signals in L1 seems to exceed the expected number relative to other elements. For instance, the base composition of the lizard L1 and the fish CR1 are almost identical (66.8 and 65.8% AT, respectively), yet there are almost three times more polyA signals in lizard L1 (23.0 ± 6.5) than in fish CR1 (8.5 ± 3.2). Similarly, the lizard Penelope has the same base composition as the mammalian L1 (62.7 and 62.3% AT, respectively), yet the mammalian L1 has on average 2.5 times more polyA (13.5 ± 3.5) than the Penelope (5.4 ± 2.6). This suggests either a stronger selection against polyA signals in non-L1 clades or selection in favor of polyA in L1, possibly to tune the level of transcription to a level tolerable by the host.

### Mutation pattern

We then investigated if the pattern of mutations in the different elements can account for the difference in base composition. To this end we estimated the relative proportion of mutations in genomic copies relative to the consensus sequence. We chose to eliminate mutations shared by multiple elements since these mutations are likely inherited from a common progenitor. Thus, we focused our attention on singletons, i.e. mutations that are unique to a sequence. Figure 11 shows the normalized proportion of *de novo* mutations for L1 and L2 in lizard, frog and fish. Some general trends are apparent, but there are also some differences among elements. In all cases, mutations from C to T and mutations from G to A are the most frequent (except for frog L1 for which we found a large proportion of T to C mutation), and this is true even when mutations in CpG are excluded. Interestingly, the strength of this mutational bias is clade specific. For L1, 42 to 46% of novel mutations are C to T or G to A while this proportion is 33 to 34% for L2. Since the lizard, frog and fish L2 have drastically different base composition, this bias is not related to the nucleotide content of the elements or its CpG content (which is very different among species). Although mutations from C to T are more common than T to C and mutations from G to A are more common than A to G (from 1.2 to 4.8-fold), there are differences among clades and among species. For instance, mutations from T to C and from A to G in lizard L2 and frog L1 account for ~30% of all mutations while they account for 14 to 19% for the other elements. Whatever the cause of these differences, it remains that all elements experience a mutation pressure toward an AT-rich nucleotide content, and when all mutation types are combined, we calculated an overall excess of mutations from GC to AT ranging from 1.4 to 2.9.

**Figure 11.**
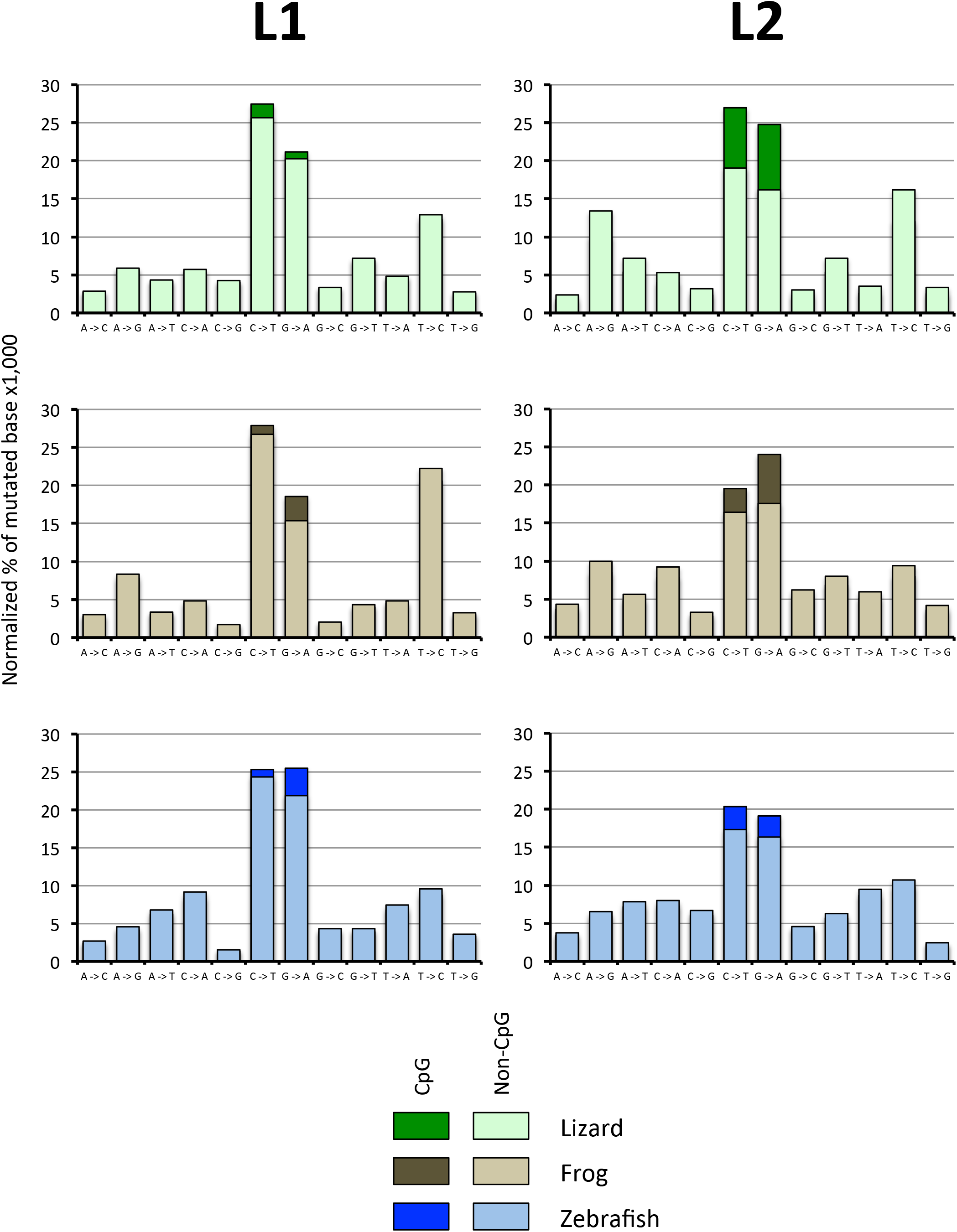
Normalized percentage of mutations in genomic copies of L1 and L2 in lizard, frog and fish. Only singletons are shown (see text).

## DISCUSSION

We performed a comprehensive analysis of the base composition of the major clades of nLTR-RTs active in vertebrates. Our results can be summarized as follows. First, we showed that the nucleotide content differs markedly among clades of nLTR-RTs within the same host and that elements belonging to the same clade can differ in base composition among hosts. Using phylogenetic analysis, we demonstrated that nucleotide content remains constant within the same host over an extended period of evolutionary time. It had been shown that the base composition of TEs differs from the base composition of hosts’ genes [38, 39] and that TEs show a tendency to be enriched in AT at the third position of codons, independently of the genome of origin [24, 40]. Based on this difference, it was proposed that the unusual nucleotide content of TEs can be used to identify them in genomes [39]. Here we showed that this picture needs to be revised and that there are some subtle differences in base composition among elements of the same clade (for instance the A-bias of L1 in mammals, lizard and frog *versus* the similar abundance of A and T in fish) but also some large differences among elements found in the same genomes (for instance L1 and L2 in lizard).

### Base composition variation and Horizontal Gene Transfer

A possible explanation for the heterogeneity in base composition within host is horizontal gene transfer (HGT). There are many cases of HGT across kingdoms of class II elements (e.g. DNA transposons) and some of these cases involve vertebrate hosts [41–43]. Such events are however exceedingly rare for nLTR-RTs. A recent investigation of HGT based on 759 eukaryotic genomes [11] showed that, although possible, HGT of nLTR-RTs remains largely limited to elements of the RTE clade [9, 44]. The same study identified six potential instances of HGT of Tx1 elements involving aquatic organisms (mostly invertebrates) but these proposed cases are extremely ancient and most cases are supported by a very small number of sequences. These authors also propose a seventh case, namely the HGT of L1 in mammals but their only argument is the apparent lack of L1 elements in monotremes and they are ignoring other arguments in favor of the vertical transmission of L1 in vertebrates, for instance the fact that the phylogeny of L1 in vertebrates matches perfectly the phylogeny of their hosts [16, 45]. Even if nLTR-RTs, other than RTE, were shown to be transferred horizontally in vertebrates, the data currently at hand support the idea that such cases would be extremely rare, and that vertical transmission is the main mode of transfer of nLTR-RT in vertebrates.

Our dataset however contains some indications that HGT has occurred and could explain some of the differences and similarities in base composition. First, we found within the same genome (Medaka) three RTE families with very different base composition, suggestive of horizontal transmissions from different sources. Second, we found a Rex1 element in frog that showed a higher level of identity with a fish element than this element had with other fish Rex1. Finally, we found in Tx1, four GC-rich sequences nested among AT-rich fish Tx1, which could be explained by HGT. These different cases will need to be analyzed in more details, however, at the time of our study, BLAST searches of public databases using those sequences did not produce any further support for HGT.

For the other clades analyzed here there is no reason to believe HGT played a role, however we can’t fully exclude it. It is indeed possible that the presence of some nLTR-RT lineages in a genome is the result of an ancient event of HGT from a host with a very different base composition. For instance, it is plausible that the GC-rich L2 element of lizard was acquired a long time ago from an unidentified host. It remains however that this element persisted and diversified in the genome of the lizard for an extended period of evolutionary time and yet retained the same base composition, despite a mutational pressure toward a higher AT content. The same reasoning can be applied to the CT-rich CR1 elements found in frogs and fish. Thus, even if we hypothesize the ancient transfer of nLTR-RT harboring different base composition, the persistence of the nucleotide content over long periods of evolutionary time remains to be explained.

### Base composition is likely maintained by selection

There are two main categories of mechanisms that can account for differences in base composition among clades of nLTR-RTs. First, if elements are exposed to different mutational processes, this can lead to different base composition over time. However, an excess of mutations toward AT is observed for all types of elements, even elements that are not AT-rich, and thus mutations alone does not appear to be the main cause of difference in nucleotide content among nLTR-RTs. This is not to say that mutation does not play a role as we did observe some differences in the pattern of mutations of elements among clades and among hosts. For instance, the lowest bias in favor of AT was found for the GC-rich L2 element of lizard and for the CT-rich L1 element of frog, and could thus contribute to the unusual base composition of these elements. The causes of the differences in mutational bias are unclear and will require further investigations that are beyond the scope of this study. Among possible factors, mutations at the hyper-mutable CpG dinucleotides, whose repair affects the probability of mutations at non-CpG sites [46–48], editing by APOBEC proteins [49], that have been shown to differentially affect L1 and L2 elements in lizard [50], or GC-biased gene conversion, which will affect differently elements that reside in regions of different recombination [51, 52] will need to be examined.

Alternatively, selective processes may dictate base composition. There are three lines of evidence suggesting that selection plays a role in the evolution of base composition. First, the base composition of a clade remains stable for extended period of evolutionary time within the same host, even though there are no intrinsic reasons why ORF2 should necessarily be AT-rich, as demonstrated by the diversity in nucleotide content of this ORF. This is exemplified in L1, which has retained an AT-rich composition with an A bias in lizard and an AT-rich composition with no A bias in fish, although L1 experienced intense lineage diversification in these organisms. Second, clades of nLTR-RTs have retained their base composition despite a mutational pressure that should have pushed them all toward an AT-rich composition, yet some clades (L2 in lizard and frog) have retained a GC-rich or CT-rich composition. This means that these mutations towards AT are not recruited in the active lineages, possibly because they are not favorable to the replicative success of the elements. Third, when there are more than one ORF the base composition of the two ORFs differs, while a strict mutational model predicts that the two ORFs should harbor similar nucleotide content. This is exemplified in L1, where the AT content in ORF2 is always higher than in ORF1.

What could be the basis of this selection? A possible explanation is that the base composition reflects selection for transcriptional efficiency or inefficiency, depending on the elements. It had been demonstrated that the AT richness of most nLTR-RTs results in poor transcription, possibly as a means of self-regulation of the elements [21, 23]. The fact that in the most AT-rich elements, the three positions of codons are similarly AT-rich is consistent with a process that is independent of the protein-coding capabilities of the ORF. The base composition of Rex1 and L2 in lizard and frog does not fit a self-regulation model since their ORFs are not enriched in AT and potential premature poly-adenylation signals. This leads to two testable hypotheses. Either these elements are transcribed at a much higher rate than AT-rich elements and are repressed (or self-regulated) by other means, or they are intrinsically less efficiently transcribed because of a weak internal promoter. These two hypotheses will require experimental testing.

It is however also likely that selection acts at the translational level. Although the synonymous codon usage does not deviate from expectation (as indicated by the CAI analysis), the overall codon usage of ORF2 is always more similar to the codon of the host than expected given the base composition of the element (as suggested by the observed RCDI values which are always lower than the expected one), which is indicative of a certain level of adaptation to the host. Selection for a more optimal codon usage could also explain why the base composition of ORF1 is less biased than ORF2, since ORF1p needs to be produced in much larger amount that ORF2p for successful transposition. It is thus likely that the base composition of nLTR-RTs is evolving in response to the joint effect of selection for lower (or higher) transcription and selection for more (or less) efficient translation of the ORFs.

### Conclusions

Our analysis on base composition evolution provides some insights on the nature of the interactions between TEs and their host and among TEs within a genome. The persistence over long time scales of base compositions that are not optimal for the replication of elements support a model of co-existence between nLTR-RTs and their hosts. Interactions between TEs and their host can range from an arms race, where hosts evolve repression mechanisms imposing a selective pressure on TEs to evade repression, to domestication, where TEs and the hosts are peacefully co-existing because they are both indispensable to the survival of each other. There are examples of both models in the literature, however it is unclear which model is the most common in nature. In a recent review, Cosby et al. examined in great details the literature on this topic and they proposed that the arms race model may not be the most prevalent one [53], but that instead, strategies that would allow TEs to persist and multiply without jeopardizing the fitness of the host have been overlooked. In this context the stability of sub-optimal base composition may provide an example of self-regulation of nLTR-RTs to maintain a harmonious relationship with their hosts. Another implication of this research is that it supports the idea that the genome is comparable to an ecosystem in which TEs compete for host resources, i.e. the community ecology of the genome [54, 55]. If we push this metaphor a little further, TEs can possibly occupy different “genomic niches” if they don’t use the same resources and thus co-exist in the genome of their hosts. The co-existence of L1 and L2 in lizard illustrates this scenario. Given their base composition, it is likely that these two elements do not use the same pool of tRNA and of amino acids for their translation and therefore occupy different niches in the genome of their host. The idea that nLTR-RTs can coexist because they differ in their use of resources will need to be better studied both theoretically and experimentally, but present an intriguing research direction to understand the mechanisms that account for the diversity of TEs in genomes.

## METHODS

The majority of the sequences analyzed here had previously been described [16, 36, 56] or were obtained from Repbase (https://www.girinst.org/repbase/). For all consensi we verified that the ORFs were intact. In the few cases they were not intact, we collected genomic copies and refined the consensus sequences. Note that we only analyzed families with a divergence from consensus lower than 5% to reduce the uncertainty in building consensus sequences. Sequences were aligned at the DNA and protein level using Geneious version 8.1.5 (www.geneious.com), which was used to estimate nucleotide content. Geneious

8.1.5 was also used to estimate the level of identity among amino acid sequences. Phylogenetic reconstructions were performed on protein sequences using the maximum likelihood method in MEGA 6.06 [57]. The robustness of the trees was assessed using 500 bootstrap resamplings.

Analyses of the base composition at the codon level were performed using the CAIcal platform at http://genomes.urv.es/CAIcal [58]. This site calculates the base composition at the three positions of codons and estimates statistics used to assess codon bias. For each codon we estimated the Relative Synonymous Codon Usage (RSCU), defined as the number of times a codon is used, divided by the number of synonymous codons encoding the same amino acid [59]. We also calculated three estimators of codon usage bias, Nc, CAI and RCDI. Nc is the effective number of codons and quantifies how much the use of a specific codon in a gene deviates from equal use of all synonymous codons [60]. Its value ranges from 20, when each amino acid is encoded by a single synonymous codon, through 61, when all synonymous codons are equally represented. The Codon Adaptation Index, CAI [61], estimates codon bias given the codon usage of an organism and the nucleotide content of the gene. The CAI ranges from 0 to 1, a value of 1 indicating that it is the most common synonymous codon that is used, which is suggestive of a low codon bias. Significance of CAI is determined by comparing the observed values of CAI with the expected CAI (eCAI), which is an estimator of the random codon usage assuming the base composition of the sequence studied [62]. Finally, we calculated the Relative Codon Deoptimization Index (RCDI), which is a measure of how different the codon usage in a gene is relative to a reference set [63]. The higher the similarity between codon usage of the host and the sequence of interest, the closer the value of RCDI is to 1. Statistical significance of RCDI was assessed by calculating the expected RCDI (eRCDI), which is determined by generating random sequences with similar nucleotide content and amino acid composition to the input sequence [64].

We determined the pattern of mutation of each clade in each species by collecting at least 8 genomic copies for each family. The alignments consisted of the entire second open-reading frame (ORF2). The genomic copies were aligned to each other and to the consensus of their respective family. The different types of mutations were tabulated using the “Find variations/SNPs” in Geneious 8.1.5. Differences among elements may result from *de novo* mutations or from differences they inherited from their progenitor. Since we were interested specifically in the type of mutations that elements experience after, or at the time of insertion, we excluded mutations that were shared between elements since those were likely inherited from a common progenitor, and may have been filtered by selective processes (due to the constraints acting on the ORF of the elements). Thus, only singletons were compared among sequences. Note that the families analyzed here are very young and most likely still active in their host. Thus, determining the ancestral and derived state of the nucleotide is trivial. For mutations from C to T and from G to A, we distinguished mutations in CpG dinucleotides from mutations in non-CpG context. This is because C and G in a CpG context mutate at a rate 10 to 50 times higher than in non-CpG [65–67].

## DECLARATIONS

### Availability of data and material

The consensus sequences analyzed here were obtained from RepBase or as supplementary material in [16, 36, 56]. The raw dataset is available as an excel spreadsheet in supplementary material.

### Ethics approval and consent to participate

Not applicable.

### Consent for publication

Not applicable.

### Author’s contribution

RR and SB generated the datasets, analyzed the data and wrote the manuscript. All authors are taking full responsibility for the work described in this article.

## ACKOWLEDGEMENTS

We thank Imtiyaz Hariyani, Yann Bourgeois, Justin Wilcox and Sebastian Kirchhof for their helpful comments on the manuscript.

## References

1. Tollis M, Boissinot S: The evolutionary dynamics of transposable elements in eukaryote genomes. Genome Dyn 2012, 7:68–91.

2. Chalopin D, Naville M, Plard F, Galiana D, Volff JN: Comparative analysis of transposable elements highlights mobilome diversity and evolution in vertebrates. Genome biology and evolution 2015, 7(2):567–580.

3. Warren IA, Naville M, Chalopin D, Levin P, Berger CS, Galiana D, Volff JN: Evolutionary impact of transposable elements on genomic diversity and lineage-specific innovation in vertebrates. Chromosome research: an international journal on the molecular, supramolecular and evolutionary aspects of chromosome biology 2015, 23(3):505–531.

4. Sotero-Caio CG, Platt RN, 2nd, Suh A, Ray DA: Evolution and Diversity of Transposable Elements in Vertebrate Genomes. Genome biology and evolution 2017, 9(1):161–177.

5. Malik HS, Burke WD, Eickbush TH: The age and evolution of non-LTR retrotransposable elements. Molecular biology and evolution 1999, 16(6):793–805.

6. Kapitonov VV, Tempel S, Jurka J: Simple and fast classification of non-LTR retrotransposons based on phylogeny of their RT domain protein sequences. Gene 2009, 448(2):207–213.

7. Luan DD, Korman MH, Jakubczak JL, Eickbush TH: Reverse transcription of R2Bm RNA is primed by a nick at the chromosomal target site: a mechanism for non-LTR retrotransposition. Cell 1993, 72(4):595–605.

8. Cost GJ, Feng Q, Jacquier A, Boeke JD: Human L1 element target-primed reverse transcription in vitro. Embo J 2002, 21(21):5899–5910.

9. Kordis D, Gubensek F: Unusual horizontal transfer of a long interspersed nuclear element between distant vertebrate classes. Proceedings of the National Academy of Sciences of the United States of America 1998, 95(18):10704–10709.

10. Gentles AJ, Wakefield MJ, Kohany O, Gu W, Batzer MA, Pollock DD, Jurka J: Evolutionary dynamics of transposable elements in the short-tailed opossum Monodelphis domestica. Genome research 2007, 17(7):992–1004.

11. Ivancevic AM, Kortschak RD, Bertozzi T, Adelson DL: Horizontal transfer of BovB and L1 retrotransposons in eukaryotes. Genome biology 2018, 19(1):85.

12. Boissinot S, Davis J, Entezam A, Petrov D, Furano AV: Fitness cost of LINE-1 (L1) activity in humans. Proceedings of the National Academy of Sciences of the United States of America 2006, 103:9590–9594.

13. Boissinot S, Entezam A, Furano AV: Selection against deleterious LINE-1-containing loci in the human lineage. Molecular biology and evolution 2001, 18(6):926–935.

14. Hancks DC, Kazazian HH, Jr.: Roles for retrotransposon insertions in human disease. Mobile DNA 2016, 7:9.

15. Goodier JL: Restricting retrotransposons: a review. Mobile DNA 2016, 7:16.

16. Boissinot S, Sookdeo A: The Evolution of LINE-1 in Vertebrates. Genome biology and evolution 2016, 8(12):3485–3507.

17. Khan H, Smit A, Boissinot S: Molecular evolution and tempo of amplification of human LINE-1 retrotransposons since the origin of primates. Genome research 2006, 16(1):78–87.

18. Boissinot S, Furano AV: Adaptive evolution in LINE-1 retrotransposons. Molecular biology and evolution 2001, 18(12):2186–2194.

19. Adey NB, Schichman SA, Graham DK, Peterson SN, Edgell MH, Hutchison CA, 3rd: Rodent L1 evolution has been driven by a single dominant lineage that has repeatedly acquired new transcriptional regulatory sequences. Molecular biology and evolution 1994, 11(5):778–789.

20. Jacobs FM, Greenberg D, Nguyen N, Haeussler M, Ewing AD, Katzman S, Paten B, Salama SR, Haussler D: An evolutionary arms race between KRAB zinc-finger genes ZNF91/93 and SVA/L1 retrotransposons. Nature 2014, 516(7530):242–245.

21. Perepelitsa-Belancio V, Deininger PL: RNA truncation by premature polyadenylation attenuates human mobile element activity. Nature genetics 2003, 35:363–366.

22. Han JS, Szak ST, Boeke JD: Transcriptional disruption by the L1 retrotransposon and implications for mammalian transcriptomes. Nature 2004, 429:268–274.

23. Han JS, Boeke JD: A highly active synthetic mammalian retrotransposon. Nature 2004, 429(6989):314–318.

24. Lerat E, Capy P, Biemont C: Codon usage by transposable elements and their host genes in five species. Journal of molecular evolution 2002, 54(5):625–637.

25. Charlesworth B, Langley CH: The evolution of self-regulated transposition of transposable elements. Genetics 1986, 112(2):359–383.

26. Zhou Z, Dang Y, Zhou M, Li L, Yu CH, Fu J, Chen S, Liu Y: Codon usage is an important determinant of gene expression levels largely through its effects on transcription. Proceedings of the National Academy of Sciences of the United States of America 2016, 113(41):E6117–E6125.

27. Plotkin JB, Kudla G: Synonymous but not the same: the causes and consequences of codon bias. Nature reviews Genetics 2011, 12(1):32–42.

28. Hershberg R, Petrov DA: Selection on codon bias. Annual review of genetics 2008, 42:287–299.

29. Sueoka N: Correlation between Base Composition of Deoxyribonucleic Acid and Amino Acid Composition of Protein. Proceedings of the National Academy of Sciences of the United States of America 1961, 47(8):1141–1149.

30. Han JS, Boeke JD: LINE-1 retrotransposons: Modulators of quantity and quality of mammalian gene expression? BioEssays: news and reviews in molecular, cellular and developmental biology 2005, 27(8):775–784.

31. Medstrand P, van de Lagemaat LN, Mager DL: Retroelement distributions in the human genome: variations associated with age and proximity to genes. Genome research 2002, 12:1483–1495.

32. Slotkin RK, Martienssen R: Transposable elements and the epigenetic regulation of the genome. Nature reviews Genetics 2007, 8(4):272–285.

33. Lerat E, Capy P, Biemont C: The relative abundance of dinucleotides in transposable elements in five species. Molecular biology and evolution 2002, 19(6):964–967.

34. Burge C, Campbell AM, Karlin S: Over- and under-representation of short oligonucleotides in DNA sequences. Proceedings of the National Academy of Sciences of the United States of America 1992, 89(4):1358–1362.

35. Furano AV, Duvernell D, Boissinot S: L1 (LINE-1) retrotransposon diversity differs dramatically between mammals and fish. Trends in genetics: TIG 2004, 20(1):9–14.

36. Novick PA, Basta H, Floumanhaft M, McClure MA, Boissinot S: The evolutionary dynamics of autonomous non-LTR retrotransposons in the lizard *Anolis carolinensis* shows more similarity to fish than mammals. Molecular biology and evolution 2009, 26(8):1811–1822.

37. Blair JE, Hedges SB: Molecular phylogeny and divergence times of deuterostome animals. Molecular biology and evolution 2005, 22(11):2275–2284.

38. Shields DC, Sharp PM: Evidence that mutation patterns vary among Drosophila transposable elements. Journal of molecular biology 1989, 207(4):843–846.

39. Andrieu O, Fiston AS, Anxolabehere D, Quesneville H: Detection of transposable elements by their compositional bias. BMC bioinformatics 2004, 5:94.

40. Jia J, Xue Q: Codon usage biases of transposable elements and host nuclear genes in Arabidopsis thaliana and Oryza sativa. Genomics, proteomics & bioinformatics 2009, 7(4):175–184.

41. Novick P, Smith J, Ray D, Boissinot S: Independent and parallel lateral transfer of DNA transposons in tetrapod genomes. Gene 2010, 449(1-2):85–94.

42. Pace JK, 2nd, Gilbert C, Clark MS, Feschotte C: Repeated horizontal transfer of a DNA transposon in mammals and other tetrapods. Proceedings of the National Academy of Sciences of the United States of America 2008, 105(44):17023–17028.

43. Schaack S, Gilbert C, Feschotte C: Promiscuous DNA: horizontal transfer of transposable elements and why it matters for eukaryotic evolution. Trends in ecology & evolution 2010, 25(9):537–546.

44. Walsh AM, Kortschak RD, Gardner MG, Bertozzi T, Adelson DL: Widespread horizontal transfer of retrotransposons. Proceedings of the National Academy of Sciences of the United States of America 2013, 110(3):1012–1016.

45. Kordis D, Lovsin N, Gubensek F: Phylogenomic analysis of the L1 retrotransposons in Deuterostomia. Systematic biology 2006, 55(6):886–901.

46. Chen J, Miller BF, Furano AV: Repair of naturally occurring mismatches can induce mutations in flanking DNA. eLife 2014, 3:e02001.

47. Walser JC, Furano AV: The mutational spectrum of non-CpG DNA varies with CpG content. Genome research 2010, 20(7):875–882.

48. Walser JC, Ponger L, Furano AV: CpG dinucleotides and the mutation rate of non-CpG DNA. Genome research 2008, 18(9):1403–1414.

49. Carmi S, Church GM, Levanon EY: Large-scale DNA editing of retrotransposons accelerates mammalian genome evolution. Nature communications 2011, 2:519.

50. Lindic N, Budic M, Petan T, Knisbacher BA, Levanon EY, Lovsin N: Differential inhibition of LINE1 and LINE2 retrotransposition by vertebrate AID/APOBEC proteins. Retrovirology 2013, 10:156.

51. Duret L, Galtier N: Biased gene conversion and the evolution of mammalian genomic landscapes. Annual review of genomics and human genetics 2009, 10:285–311.

52. Mugal CF, Weber CC, Ellegren H: GC-biased gene conversion links the recombination landscape and demography to genomic base composition: GC-biased gene conversion drives genomic base composition across a wide range of species. BioEssays: news and reviews in molecular, cellular and developmental biology 2015, 37(12):1317–1326.

53. Cosby RL, Chang NC, Feschotte C: Host-transposon interactions: conflict, cooperation, and cooption. Genes & development 2019, 33(17-18):1098–1116.

54. Brookfield JF: The ecology of the genome - mobile DNA elements and their hosts. Nature reviews Genetics 2005, 6(2):128–136.

55. Venner S, Feschotte C, Biemont C: Dynamics of transposable elements: towards a community ecology of the genome. Trends in genetics: TIG 2009, 25(7):317–323.

56. Blass E, Bell M, Boissinot S: Accumulation and rapid decay of non-LTR retrotransposons in the genome of the three-spine stickleback. Genome biology and evolution 2012, 4(5):687–702.

57. Tamura K, Peterson D, Peterson N, Stecher G, Nei M, Kumar S: MEGA5: Molecular Evolutionary Genetics Analysis using Maximum Likelihood, Evolutionary Distance and Maximum Parsimony Methods. Molecular biology and evolution 2011, 28:2731–2739.

58. Puigbo P, Bravo IG, Garcia-Vallve S: CAIcal: a combined set of tools to assess codon usage adaptation. Biology direct 2008, 3:38.

59. Sharp PM, Tuohy TM, Mosurski KR: Codon usage in yeast: cluster analysis clearly differentiates highly and lowly expressed genes. Nucleic acids research 1986, 14(13):5125–5143.

60. Wright F: The ‘effective number of codons’ used in a gene. Gene 1990, 87(1):23–29.

61. Sharp PM, Li WH: The codon Adaptation Index--a measure of directional synonymous codon usage bias, and its potential applications. Nucleic acids research 1987, 15(3):1281–1295.

62. Puigbo P, Bravo IG, Garcia-Vallve S: E-CAI: a novel server to estimate an expected value of Codon Adaptation Index (eCAI). BMC bioinformatics 2008, 9:65.

63. Mueller S, Papamichail D, Coleman JR, Skiena S, Wimmer E: Reduction of the rate of poliovirus protein synthesis through large-scale codon deoptimization causes attenuation of viral virulence by lowering specific infectivity. Journal of virology 2006, 80(19):9687–9696.

64. Puigbo P, Aragones L, Garcia-Vallve S: RCDI/eRCDI: a web-server to estimate codon usage deoptimization. BMC research notes 2010, 3:87.

65. Duncan BK, Miller JH: Mutagenic deamination of cytosine residues in DNA. Nature 1980, 287(5782):560–561.

66. Sved J, Bird A: The expected equilibrium of the CpG dinucleotide in vertebrate genomes under a mutation model. Proceedings of the National Academy of Sciences of the United States of America 1990, 87(12):4692–4696.

67. Hwang DG, Green P: Bayesian Markov chain Monte Carlo sequence analysis reveals varying neutral substitution patterns in mammalian evolution. Proceedings of the National Academy of Sciences of the United States of America 2004, 101(39):13994–14001.

